# Model-guided design of the diversity of a synthetic human gut community

**DOI:** 10.1101/2022.03.14.484355

**Authors:** Bryce M. Connors, Sarah Ertmer, Ryan L. Clark, Jaron Thompson, Brian F. Pfleger, Ophelia S. Venturelli

## Abstract

Microbial communities have tremendous potential as therapeutics. However, a major bottleneck is manufacturing high-diversity microbial communities with desired species compositions. We develop a two-stage, model-guided framework to produce microbial communities with target species compositions. We apply this method to optimize the diversity of a synthetic human gut community. The first stage exploits media components to enable uniform growth responses of individual species and the second stage uses a design-test-learn cycle with initial species abundance as a control point to manipulate community composition. Our designed culture conditions yield 91% of the maximum possible diversity. Leveraging these data, we construct a dynamic ecological model to guide the design of lower-order communities with desired temporal properties over a longer timescale. In sum, a deeper understanding of how microbial community assembly responds to changes in environmental factors, initial species abundances, and inter-species interactions can enable the predictable design of community dynamics.

## INTRODUCTION

The potential of microbial communities as human therapeutics is evidenced by the remarkable efficacy of fecal microbiota transplantation (FMT) in treating recurrent *C. difficile* infection^1^. This strategy of modifying a patient’s dysbiotic microbiome with live, therapeutic organisms (“bugs-as-drugs”) holds significant promise for treating an ever-lengthening list of microbiome associated health conditions^2^. However, FMT also poses the risk of pathogen transmission and other adverse health outcomes^3–5^. Further difficulties with this procedure include development of regulatory standards, definition of a precise mechanism of action, and scalability of donor material supply chain^6,7^. A promising alternative is the use of defined microbial community therapeutics^8^. The beneficial properties of these well-characterized mixtures of isolates could be optimized while avoiding the drawbacks of FMT^9–11^.

A key challenge towards this goal is the scalable production of defined, therapeutic communities that span the phylogenetic and functional diversity of the healthy adult microbiome^12^. Most of the commercially successful “probiotics” that are commonly recommended by physicians have gained traction not because of conclusive clinical indications, but rather because they are relatively easy to produce^13,14^. “Probiotics” tend to be oxygen-tolerant anaerobes like *Lactobacilli* and *Bifidobacterium*, while the healthy adult microbiome tends to be dominated by fastidious, oxygen-sensitive anaerobes such as *Bacteroides, Prevotella, Clostridiaceae, Ruminococcaceae, and Lachnospiraceae*^15^. “Probiotics” have even been shown to impair post-antibiotic microbiome recovery^16^. The challenge of producing therapeutic communities is a barrier to more than just commercial manufacturing; it slows scientific progress by limiting pilot-scale drug supply to clinical trials and precludes low-cost, global health applications^13,17,18^. A major contribution to this production challenge is the current strain culturing process, in which the constituent organisms of the community are grown as separate cultures, then subsequently mixed to a desired species composition^18^. This process is complicated, costly, and scales poorly for communities with large numbers of organisms^18^. Therefore, new methods to produce microbial communities with desired species compositions could alleviate this manufacturing bottleneck.

Developing model-guided approaches to predict community growth as a function of specific control inputs would greatly enhance our ability to manipulate community composition towards a desired state^19^. Design of experiments with statistical modeling (DoE) has been increasingly used to study and engineer biological systems. For example, DoE has been used to explore regulatory sequence space for modulating protein translation and for tuning enzyme expression to optimize production of a target metabolite^20–22^. In addition, DoE was used to design chemically defined media by optimizing microbial growth as a function of various media components^23,24^. Statistical modeling, an integral part of the DoE workflow, has been applied to predict microbial community composition as a function of dietary inputs, though it has been more commonly used to predict a given community-level function from species abundance^25–27^. Dynamic ecological models, while generally lacking abiotic control points like resources, have been shown to be predictive of microbial community assembly in a particular media environment^28,29^. These studies have demonstrated that inter-species interactions and initial species abundances strongly affect transient states of community assembly, suggesting that these parameters could be used to manipulate community dynamics.

We develop a two-stage, model-guided approach for systematically tuning key media components and initial species densities to optimize the diversity of a synthetic human gut community. Using statistical modeling, we design a new culture medium that yields a more uniform distribution of endpoint abundances of the monocultures. This monoculture-based optimization procedure improves community diversity. Then, in communities cultured on the new medium, we use a design-test-learn cycle to modulate individual species’ initial population densities (i.e., inocula) to further optimize community diversity. In both stages, a substantial degree of community composition (a systems-level property) can be forecasted as the composite behavior of constituent monocultures (parts-level properties)^30^. Finally, we use our data to build a dynamic ecological model that captures inter-species interactions and use this model to guide the design of communities with distinct classes of dynamic behaviors. In sum, we demonstrate that model-guided design of experiments can be combined with high-throughput species abundance measurements to steer community composition towards desired states.

### Manipulating media components to enhance community Shannon diversity

The diversity of a donor’s microbiota has been identified as a major factor determining clinical response during the use of FMT to treat inflammatory bowel disease^31,32^. Since diverse, defined communities are useful therapeutics, we aimed to maximize the Shannon diversity (Methods, equation 1) of a synthetic human gut community^10,11^. Shannon diversity is an ecological metric used to characterize both the number of species in a community and the evenness of their population sizes^33^. We designed a representative synthetic 10-member community that spans the phylogenetic and metabolic diversity of the human gut microbiome (**Fig. 1a**). This community consisted of *Blautia hydrogenotrophica* (BH), *Bifidobacterium longum* (BL), *Bacteroides uniformis* (BU), *Collinsella aerofaciens* (CA), *Dorea longicatena* (DL), *Eggerthella lenta* (EL), *Eubacterium rectale* (ER), *Faecalibacterium prausnitzii* (FP), *Prevotella copri* (PC), and *Parabacteroides johnsonii* (PJ). Several of these species, including FP, have been shown to be critical to the recovery of a healthy microbiome after childhood malnutrition and thus hold promise as bacterial therapeutics for global health applications^34,35^.

**Figure 1.**
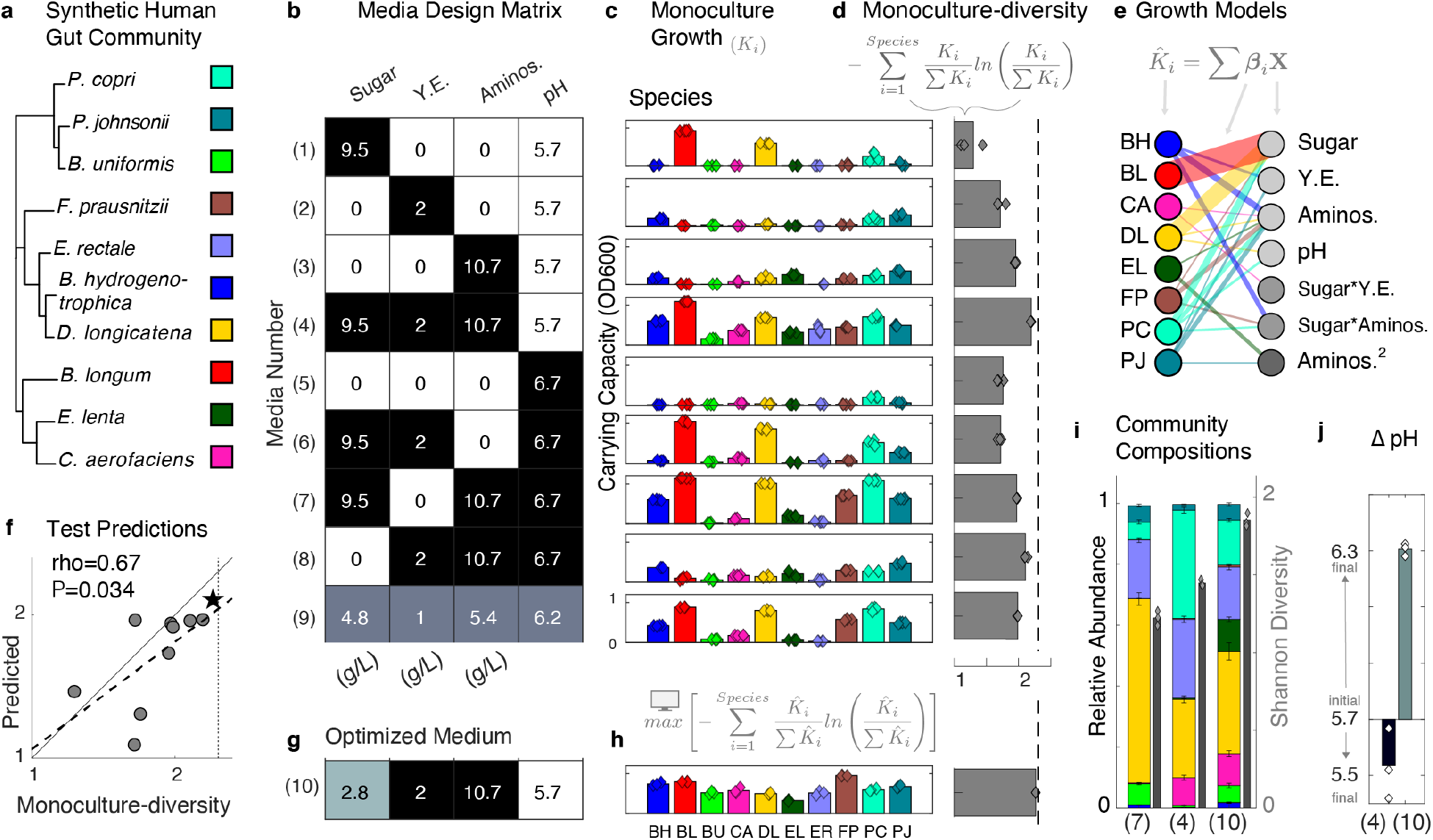
Model-guided design of media composition to enhance community Shannon diversity. **a** Phylogenetic tree of the 10-member synthetic human gut community: Bacteroidetes (upper branch), Firmicutes (middle branch), and Actinobacteria (lower branch). Phylogenetic analysis was performed using a concatenated alignment of 37 single-copy marker genes in Phylosift^42^. **b** Media factor experimental design that varies the concentration of a sugar mixture (glucose, arabinose, and maltose), yeast extract (Y.E.), defined amino acid mixture (Aminos.), and pH in a common base medium (Methods). Shading indicates design levels: “high” (black), “intermediate” (gray), and “low” (white), with concentration values labeled in units of g/L or pH. **c** Bar plots of the steady-state abundance (carrying capacity, *K_i_*) of each species determined by fitting a logistic differential equation model to the time-series measurements of absorbance at 600 nm (OD600) in each media condition (Methods, equation 3, **Fig. S1**). Different colors denote species shown in g. Bar height denotes the mean carrying capacity and data points denote biological replicates (n=4 with outlier detection, Methods). **c** Bar plots of “monoculture-diversity” (Methods, equation 4,5) based on the mean carrying capacities for each medium (Methods). Data points denote monoculture-diversities calculated from each biological replicate. Dashed line indicates maximum possible monoculture-diversity for ten species. **e** Bipartite network representation of linear regression growth models (MR, Table S1), where edge thickness is scaled by mean parameter value across cross validated parameter sets. Models predict the carrying capacity of each species 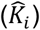 as a function of media component concentrations (**X**) (Methods, equation 6), and as such, the parameters (*β_i_*) represent the inferred growth effect of a media component on a particular species. Left and right nodes denote species and media components, respectively. Light gray nodes denote main effects, medium gray nodes denote interactions, and dark gray nodes denote quadratic main effects. Parameters with mean values of less than 0.05 are not shown. **f** Scatter plot of monoculture diversity calculated from fitted carrying capacities (x-axis) vs. monoculture-diversity calculated from media regression model validation/test predictions (“predicted”, y-axis, Methods). Pearson correlation (rho) and p-value (P) are indicated. Star indicates optimized medium. **g** Heatmap of media component concentrations that maximized monoculture-diversity (Methods, equation 7, Methods). Color scale is according to (a). **h** Bar plot of the inferred carrying capacities based on the logistic model of each species on the optimized medium (**Fig. S1b**) (f). **i** Stacked bar plot (left bars) of community compositions from the even inoculum proportion in the baseline medium (7), the highest monoculture-diversity screened medium (4), and the optimized medium (10). Bar height indicates mean of 3 biological replicates, error bars indicate 1 s.d., and all replicates are shown in Fig. S14. Shannon diversity of mean community composition (n=3 biological replicates, Methods, equation 1) is indicated as gray solid bars (right bars). Shannon diversities as calculated from each set of biological replicates are overlaid as diamonds. **j** Bar plot of the change in media pH for community cultures in the best screened (4) and optimized medium (10). Bar height indicates mean of biological replicates (diamonds, n=3).

We characterized the growth of individual species (monocultures) in a baseline defined medium that can support the growth of diverse human gut species (Methods, Supplemental Data 1)^25^. The monocultures displayed a wide range of growth rates and population sizes at steady-state (i.e. carrying capacities) (**Fig. S1a**, medium 7), suggesting that the species with low monoculture fitness may be outcompeted in the community. Human gut anaerobes have diverse metabolic strategies^36,37^. Therefore, we exploited the concentrations of key media components to manipulate monoculture growth responses^36,38^. Sugars and amino acids represented the main fermentable substrates, consistent with their key role in the mammalian gut^39^. Likewise, pH is a major environmental factor, and can distinctly modify bacterial growth^40^. In addition, we selected yeast extract since it consists of a complex digest containing vitamins, peptides, and other resources, and supports the growth of FP^41^. We used statistical design of experiments (DoE) to identify an optimal concentration profile of these components by manipulating four key variables: (1) a mixture of three sugars, (2) a defined mixture of amino acids, (3) yeast extract, and (4) pH. The “DoE” workflow involves (1) identification of (independent) variables and (dependent) responses of the system, (2) construction of an experimental design matrix of combinations of levels of each variable that satisfies a designated optimality criterion, (3) experimental implementation, (4) statistical modeling of the experimental data, and (5) use of optimization techniques to determine the values of the variables that are predicted to yield a desired system response.

We use this workflow to maximize the similarity between steady-state population sizes (i.e. carrying capacities) of the monocultures as a function of media component concentrations, while also supporting sufficient growth (**Fig. 1b**, Methods).^43^

We performed time-series measurements of optical density at 600 nm (OD600) for each monoculture in each media condition (**Fig. 1b**) and fit a logistic growth model (LM, **Table S1**) to these data (**Fig. S1a**). The carrying capacity parameter (*K_i_*) of this model indicates population size at steady-state (**Fig. 1c**). To quantify the similarity among the growth responses of individual species as a function of media components, we determined the Shannon diversity of the normalized carrying capacities in a particular medium. Normalization was performed by dividing by the sum of the inferred carrying capacities in a particular medium, mirroring how Shannon diversity is calculated from community absolute abundance data (Methods, equation 4). This quantity, hereafter referred to as “monoculture-diversity,” varied widely as a function of media composition (**Fig. 1d**).

Although we identified a medium that enabled high monoculture-diversity in the screening experiment (**Fig. 1d**, medium 4), we used model-guided optimization for further improvement. We fit linear regression models (MR, **Table S1**) with quadratic and interaction terms to predict the carrying capacity of each species from the concentrations of the media component variables (**Fig. 1e**, Methods, equation 6). The media regression model parameters provide an interpretable relationship between the concentration of a given media component and its effect on the growth of a given organism. For instance, the main effects regression parameter corresponding to “sugars” was large for the BL and DL growth models, consistent with their substantial growth improvement in the presence of the sugar mixture (**Figs. 1b,c,e**, **S2b,c**). Interaction parameters in the regression models captured more subtle trends, as these terms indicate a specific combination of independent variables that results in a distinct effect on the measured response. For example, BH had a substantial growth improvement in media containing both amino acids and sugars (**Fig. 1b,c**). The large magnitude of this interaction parameter for BH suggested that the simultaneous presence of amino acids and sugars enhanced growth more than the sum of their individual contributions alone (**Figs. 1e**, **S2a**).

To reduce overfitting and biasing of hyperparameters, we implemented elastic net regularization with nested leave-one-out cross validation (Methods). Goodness of fit was high for all species, while validation predictions on the out-of-fold measurements ranged in accuracy (**Fig. S3a**). Despite the sparse sampling of the design space using the DoE approach (**Fig. S4**), the models were predictive of an aggregate property (monoculture-diversity) on new data, even though they were variably predictive of the constituent species (**Fig. 1f**, Pearson rho=0.67, P = 0.034).

An optimization procedure (Methods, equation 7) identified a profile of media factor concentrations that maximized the predicted monoculture-diversity (Methods). The predicted concentrations were similar to medium 4, but contained 3-fold less sugar (**Fig. 1b,g**). To test this prediction, individual species were grown in the optimized medium. The monoculture-diversity for the optimized medium was close to the maximum possible value, consistent with the model prediction (**Fig. 1f,g**).

To determine if monoculture-diversity could inform the Shannon diversity of the community, we cultured the 10-member community from even initial species proportions in the baseline medium 7, best screened medium 4, and optimized medium (**Fig. 1b,g**). The model-guided, monoculture-based optimization process yielded a concomitant improvement in community Shannon diversity (**Fig. 1h,i**). Our results suggest that the reduced sugar concentration in the optimized medium, as compared with the best screened medium 4, mitigated rapid production of high levels of inhibitory organic acids by fast growing sugar fermenters. This was consistent with the substantially higher endpoint pH of a community cultured in the optimized medium 10, compared to the acidified environment of medium 4 (**Fig. 1j**). A microbial community culture that autonomously maintains non-inhibitory pH levels could be produced in simple vessels (e.g. flasks or tanks), obviating the need for expensive equipment (e.g., bioreactors with pH control).

Our model-guided, high-throughput, monoculture-based approach identified a single medium in which all species were capable of similar endpoint growth. As compared to the baseline medium, Shannon diversity of the community was increased from 53% to 80% of its maximum possible value. These results demonstrated that a moderate number of media components are effective control points for manipulating community composition.

### A constrained system of logistic equations predicts trends in community assembly

The initial population density of the constituent members of a microbial community has been shown to impact community assembly^28,44,45^. Therefore, we reasoned that we could use a design of experiments approach to further optimize community diversity as a function of inoculum density. However, this constituted a large design space for community experiments, as there are many possible combinations of inoculum proportions for a 10-member community. We first studied the “parts” of our microbial community by characterizing growth kinetics of the monocultures across a wide range of inoculum densities.

Lower inoculum density delayed the time at which the species entered a measurable exponential growth phase (**Fig. 2b**). In addition, BH, ER, and FP tended not to grow (or displayed variable growth between biological replicates) at lower inoculum densities. The remaining species displayed consistent growth kinetics at most inoculum densities, which spanned several orders of magnitude. A logistic model (LI, **Table S1**) with a single parameter set represented each species growth kinetics across the large range of inoculum densities (**Fig. 2b, Methods**).

**Figure 2.**
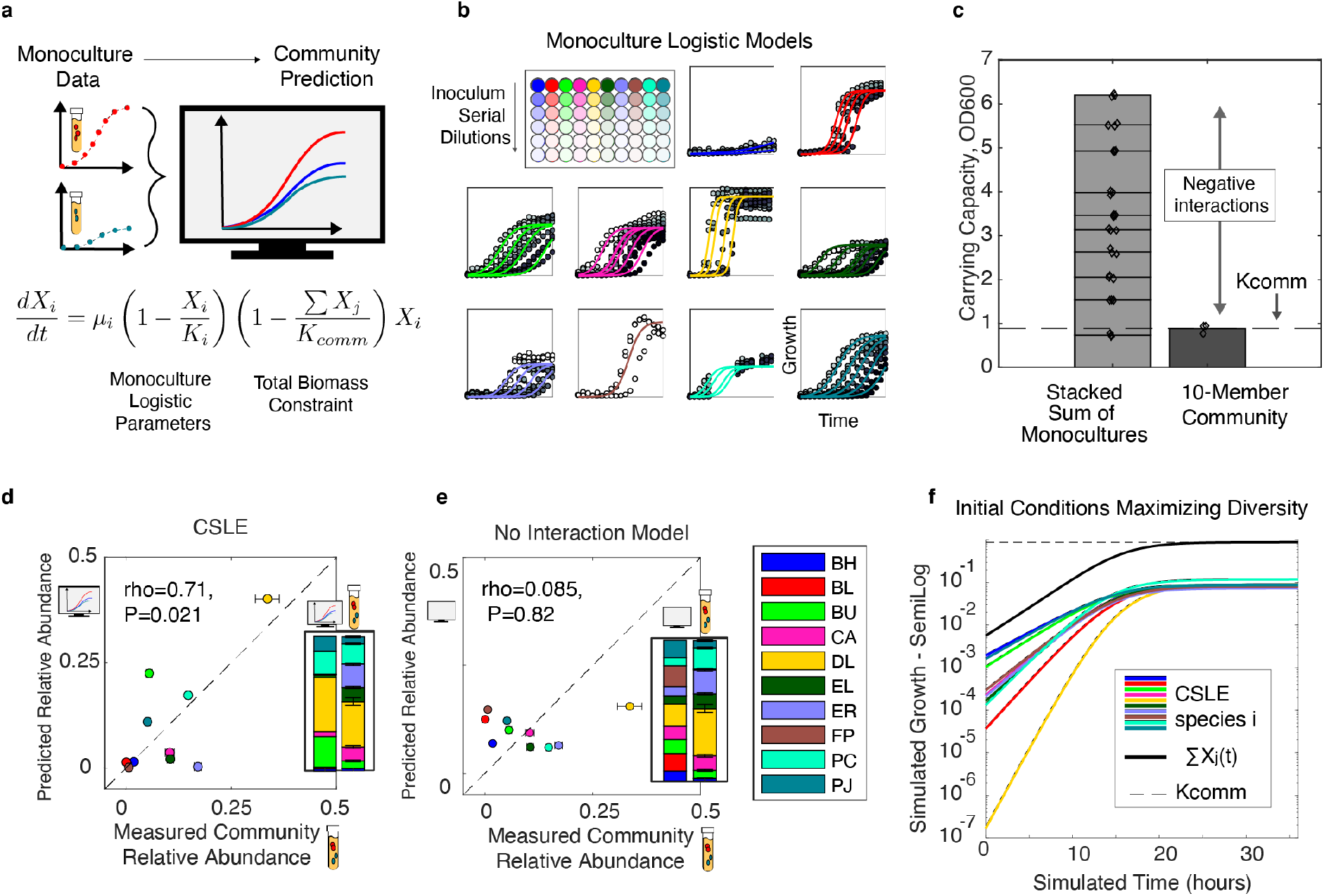
Predicting community assembly using a constrained system of logistic equations. **a** Schematic of experimental approach and model equation to predict community assembly as a set of monoculture logistic models (LI, **Table S1**) coupled via a total community growth limit, referred to as a “constrained system of logistic equations” or “CSLE” (Methods, equation 9 and Supplementary Information). Parameters: monoculture logistic growth rates (*μ_i_*), monoculture carrying capacities (*K_i_*), and total community growth limit (*K_comm_*, community carrying capacity). **b** Monoculture kinetic data based on absorbance at 600 nm (OD600, filled circles) where each species was inoculated at a range of initial densities (0.01 to 1e-7 OD600 by 10-fold serial dilution). Inoculum densities that did not yield reproducible growth were omitted (**Fig. S5a**). Lines denote the logistic differential equation fit to the time-series OD600 measurements (Methods). Colors denote species per legend in (e). **c** Bar plot of the endpoint growth of a 10-member community culture vs. the sum of the inferred logistic carrying capacities of all 10 monocultures (bar height indicates mean, diamonds show biological replicates, n=3). *K_comm_* (denoted by dashed line) represents the mean of the endpoint OD600 of the 10-member community culture (n=3 biological replicates). **d** Scatter plot of the CSLE model predictions (y-axis, left stacked bar) versus the experimentally measured community relative abundance data (x-axis, right stacked bar). Pearson correlation coefficient and p-value are indicated by “rho” and “P”, respectively. Dashed “x=y” line represents where predictions from a perfectly accurate model would fall. Error bars on experimental data denote 1 s.d. of biological replicates (n=3). **e** Scatter plot of predicted community composition based on a set of independent, logistic differential equations (y-axis, right stacked bar) and measured community composition (relative abundance, x-axis, lefthand stacked bar). Pearson correlation coefficient and p-value are indicated by “rho” and “P”, respectively. Dashed “x=y” line represents where predictions from a perfectly accurate model would fall. Error bars on experimental data denote 1 s.d. from the mean of biological replicates (n=3). **f** Line plot of CSLE simulation of monospecies growth. Optimization techniques are used to maximize the predicted Shannon diversity as a function of initial conditions (Methods, equation 10). This set of initial conditions is later used as a reference point to guide community experimental design (**Fig. 3**). Colors denote species per legend in (e).

The 10-member community cultured from an even species inoculum displayed a substantially lower total growth than the sum of the monoculture carrying capacities (**Fig. 2c**). This implies that negative inter-species interactions dominated the ecological network of the community. The total growth of microbial communities in batch culture was shown to be a saturating function of the number of species in the community^25^. Therefore, we reasoned that an upper limit on total community growth (independent of species composition) could serve as a useful null-hypothesis governing community assembly, given unknown, but largely negative, inter-species interactions. Further, we assumed that a species with higher fitness in monoculture would display higher abundance in the community.

We captured these behaviors by deriving a mathematical model, referred to as a “constrained system of logistic equations” (CSLE) (**Supplementary Information**). In this model, a species grows according to its monoculture logistic kinetics until total growth is constrained by a “community carrying capacity” (*K_comm_*). Thus, a species may cease to grow (*dx_i_/dt* → 0, arrow represents approaches) either when its population size approaches its monoculture logistic carrying capacity (*x_i_*(*t*) → *K_i_*) or when the total community growth approaches the community carrying capacity (∑*x_j_* (*t*) → *K_comm_*). *K_comm_* was defined as the mean OD600 of biological replicates of the 10-member community culture (**Fig. 2c**).

The CSLE model captured major trends in measured relative species abundances of the community (Pearson rho=0.71, P=0.021, **Fig. 2d**). Conversely, predicting community assembly as a set of independent logistic models (assuming no inter-species interactions) failed to describe community composition (Pearson rho=0.085, P=0.82, **Fig. 2e**). In the CSLE model, species that grow faster in monoculture are more likely to negatively impact the growth of other community members, resulting in a trade-off in the species’ endpoint abundances. For example, the CSLE model accurately predicted that the species with the highest monoculture growth rate (DL, yellow) would occupy a substantially larger fraction of the community than the other species (**Figs. 2d,e, S1c**). However, the set of independent logistic models failed to predict this trend. This demonstrates that the CSLE model, which was not informed by community data, could predict trends in community assembly.

The CSLE model reaches equilibrium for any community composition in which species’ absolute abundances sum to the total growth limit (*∑x_i_*(*t*) = *K_comm_*). Thus, in contrast to the logistic model, which has a single positive steady-state, the steady-state population size of a species in the CSLE model is a continuous function of initial conditions (as long as *∑K_i_ > K_comm_*, i.e., in a large community). This model allowed us to computationally explore how community composition changes as a function of species inoculum prior to collecting community data (**Fig. 2f**).

### Tuning species inoculum densities to optimize community Shannon diversity

To further optimize the endpoint Shannon diversity of the 10-member community, we used a model-guided design-test-learn (DTL) cycle to modulate the inoculum densities of each species (**Fig. 3a**). The iterative DTL approach uses models, trained on community composition data collected in previous cycles, to guide the design of experimental conditions for the subsequent cycle^25^. The “design” step was initiated with the construction of an experimental design matrix. Inoculum density values were assigned to the levels of the matrix using model predictions when possible. The “test” step used automated liquid handling to array the designed inocula conditions (Methods). Community cultures were grown to approximately stationary phase, and species abundances were analyzed using multiplexed NGS (Methods). The “learn” step inferred parameters from experimental data and evaluated the predictive capability of the statistical models.

**Figure 3.**
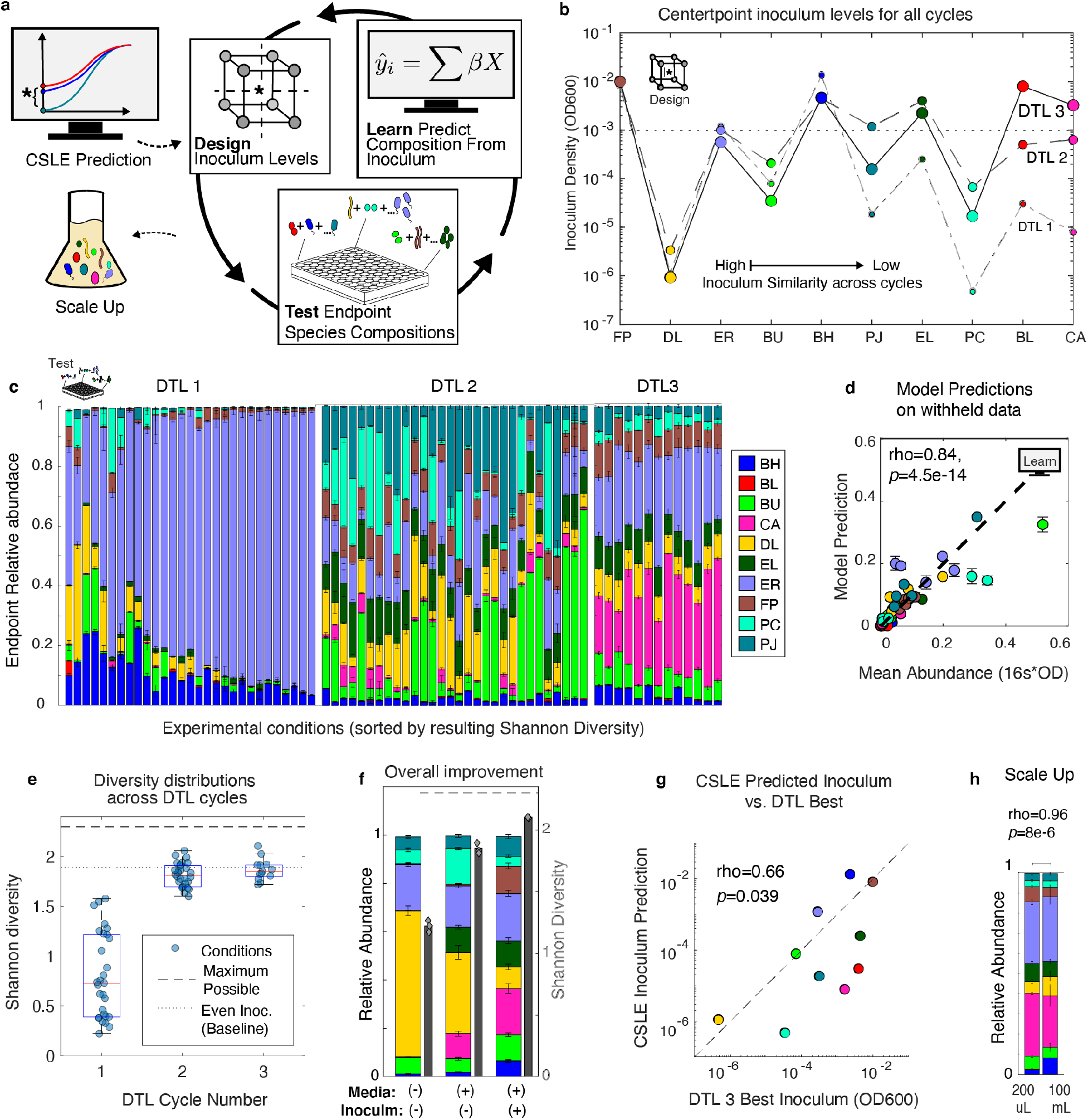
Tuning species inoculum densities to optimize community Shannon diversity in a design-test-learn cycle. **a** Schematic illustrating the design-test-learn (DTL) cycle for maximizing community diversity as a function of species inocula. The “center point” of each experimental design corresponds to the inoculum (colored circles) predicted to yield the highest community Shannon diversity. DTL 1 center point is predicted with the CSLE model, thereby exploiting monoculture growth data to design the first community experiment. Subsequent DTL cycle center points are predicted according to inoculum regression models (IR, **Table S1**), which are trained on community data collected during the DTL process. Scale-up of the culture is performed for potential bioprocessing applications (bottom left). **b** Categorial scatter plot of center point inoculum conditions for each DTL cycle, which are informed by model predictions when possible (Methods). For visualization, species are sorted by the magnitude of the difference between the log transformed inoculum densities of the first and last DTL cycles. The dotted line indicates the even inoculum baseline condition. Full design matrices are shown in Supplementary Data 3. **c** Stacked bar plots of endpoint community compositions for each DTL cycle, sorted left to right by community Shannon diversity. Stacked bars and error bars represent mean and 1 s.d. of the mean of biological replicates (n=3), respectively, for each condition; all replicates are shown in Fig. S13. **d** Scatter plot of the experimentally measured absolute abundance of each species versus the linear regression models’ predictions of endpoint species absolute abundance on the test set. The model was trained on community composition measurements from the first two DTL cycles. The dependent variable is the endpoint species abundance, and the independent variables are the initial OD600 of each species from the inoculum design matrix. Pearson correlation coefficient (rho), and p-value (P) are shown. The validation (out-of-fold) predictions with species-specific correlation coefficients are shown in **Fig S6**. **e** Distributions of Shannon diversities calculated from the mean composition of biological replicates (n=3) for conditions of each DTL cycle (blue circles). Red line in each box denotes the median, upper and lower edges denote 75^th^ and 25^th^ percentiles, respectively, and whiskers denote range of non-outlier datapoints. Dashed line indicates maximum possible Shannon diversity for a 10-member community. Dotted line indicates the diversity from even inoculum in the optimized medium (**Fig. 1i**). **f** Stacked bar plot of the community composition from media optimization (**Fig. 1**) and inoculum optimization (**Fig. 3**). Species composition (stacked bars, left-axis) and Shannon diversities as calculated from mean of species abundances (gray solid bar); diamonds show diversities calculated from individual sets of biological replicates (right-axis), and all biological replicates are shown in Fig. S14. Even inoculum and baseline medium (pre-optimization) are indicated with (-), while (+) indicates that the community resulted from media or inoculum optimization in this study. **g** Scatter plot of the log transform of inoculum densities predicted by the CSLE model (DTL1 center point levels) vs. the experimentally identified best inoculum (condition yielding highest diversity after three DTL cycles). Pearson correlation is calculated between the logarithm of the inoculum densities. **h** Stacked bar plot of species relative abundance of the 10-member community cultured in a 200 uL microtiter plate versus a 100 mL flask, bar height and error bars represent mean and 1 s.d. of 3 biological replicates, all biological replicates are plotted in Fig. S14.

The assignment of inoculum density values to the levels of the DoE design matrix for the first community inoculum experiment was guided by the CSLE model (**Fig. 2a,d**). We used optimization to solve for a set of initial conditions that maximized the predicted Shannon diversity (**Fig. 2f**). This set of initial conditions was used as a central reference point (“center-point”), representing the “medium” level for all species, around which the rest of the experimental design was constructed (**Fig. 3a**). These designs were constructed for the dual purpose of identifying a high diversity condition and collecting structured training data to improve the model’s predictive ability.

Community compositions varied widely as a function of the experimental design conditions (**Fig. 3c,** DTL 1), confirming that inoculum density was a useful control point for manipulating community assembly. Despite a modest monoculture growth rate and carrying capacity (**Fig. 2b, S1c**), ER overgrew in many conditions (**Fig. 3c**, light purple). The CSLE model had underpredicted ER in the test community grown from an even inoculum (**Fig. 2d**), suggesting that ER benefits from positive inter-species interactions that were not captured in the CSLE model.

Consequently, the CSLE model overpredicted the initial density of ER, which in turn resulted in overgrowth in the community.

This community data to was leveraged to quantify inter-species interactions beyond global competition. Regression models with linear, quadratic, and interaction terms (IR1, **Table S1**) were trained to predict the absolute abundance of each species in the community from the inoculum values of the experimental design (Methods). After the first DTL cycle, the inoculum regression models accurately predicted half of the species (Pearson rho > 0.7, P < 1e-6, **Fig. S7a**). However, three species (CA, EL and PC) with predictive models displayed low overall growth (average relative abundance less than 2.5% across design conditions, **Fig. 3c** “DTL1”,). As such, these models were not practically useful, since predicting maximum diversity (i.e., 10% relative abundance) would result in significant extrapolation of the model.

In DTL 2, the new center point inoculum value for species that were poorly predicted or displayed low overall growth was qualitatively determined based on community data. If a species tended to overgrow (ER) in the previous cycle, the new center point value was set at the previous cycle’s low value. By contrast, if a species tended to undergrow (BL, CA, DL, EL, PC and PJ), its new center point value was set to the previous high value (**Fig. S8**, Methods). We performed model-guided optimization of the inoculum values to maximize the predicted Shannon diversity for species that were accurately predicted by the model and displayed substantial growth (Methods).

These optimized values were used as the center point values for DTL 2 (Methods). In DTL 2, most species were well-represented, and ER was present at a lower abundance in the community than in DTL 1 **(Fig. 3c)**. The median community diversity was substantially higher in DTL 2 than DTL 1, indicating that data obtained in DTL 1 were informative for enhancing Shannon diversity (**Fig. 3e**). Inoculum regression models were re-trained on community composition data from both DTL 1 and 2 (IR2, **Table S1**), and the models accurately predicted the absolute abundance of all species except BL during cross validation (Pearson rho > 0.70, P < 1e-8, **S7b**). Further, the model accurately predicted test communities that were withheld from the training and validation process (**Fig. 3d**, Pearson rho=0.84, P=2.5e-14).

To determine if the Shannon diversity could be enhanced further, we used optimization techniques using the nine predictive regression models to determine a new inoculum center point for DTL 3. The high and low levels probed a smaller design space than previous cycles reflecting higher confidence based on the substantial improvement in Shannon diversity in DTL 2. Since BL consistently undergrew and was poorly predicted, its inoculum density was set to a maximum designated value (Methods). Despite having the largest inoculum and high monoculture fitness, BL was low abundance in DTL 3, indicating that BL was inhibited by other members of the community (**Fig. 3c**). Notably, the beneficial species FP was higher abundance in DTL 3 communities than in the community inoculated with an even inoculum (**Fig. 3f**)^46–48^. Overall, the highest Shannon diversity condition was identified in DTL 3, representing 91% of the maximum possible value for a 10-member community (**Fig. 3e,f**). This was a substantial improvement from the already high 80% of the maximum diversity achieved by medium optimization alone (**Fig. 3f**).

The set of inoculum densities that yielded the highest Shannon diversity in DTL 3 was correlated to the CLSE optimized inoculum prediction (Pearson rho between logarithm of inoculum values = 0.66, P = 0.039, **Fig. 3g**). Further, for half of the species, inocula for the highest diversity condition were within three-fold of the CSLE predicted values (**Figs. 2f, 3b**). These data show the CSLE model prediction was a useful starting point for the DTL cycle, as it substantially narrowed the inoculum design space that yielded assembly of a highly diverse community.

Biomanufacturing of microbial communities in a real-world setting would require (1) robustness of endpoint community composition to technical variability in species inocula, (2) translation to production-scale equipment, and (3) viability of organisms harvested at the endpoint. Despite the four-fold variation in inoculum in DTL 3, the coefficient of variation of the endpoint Shannon diversity across design conditions was less than 6% (**Fig. 3e**). This demonstrates that our process was robust to variation in species inocula. The community compositions in 200 uL and 100 mL batch cultures were similar, demonstrating that a 500-fold difference in batch culture scale did not substantially alter community assembly (**Fig. 3h**, Pearson rho=0.96, P=8e-6). To evaluate the viability of species in the endpoint community cultures, we transferred a small aliquot (25-fold volume/volume dilution) of the communities measured at the endpoint into fresh media and grew them to approximately stationary phase (**Methods**). All species in all conditions yielded greater than three-fold increase in absolute abundance during the second passage, demonstrating that these species were viable (**Fig. S9f**).

### Model-guided design of microbial community dynamics

Positive and negative inter-species interactions are major determinants of microbial community assembly^25,49^. Therefore, we constructed a dynamic ecological model that captured specific inter-species interactions (**Fig. 2d,e**). The generalized Lotka-Volterra (gLV) model (Methods, equation 13) is a set of coupled ordinary differential equations that describes a specie’s growth dynamics as a function of its basal growth rate and interactions with each constituent community member. This model has accurately predicted complex community dynamics, and its interpretable parameters have revealed significant inter-species interactions^25,29,49^.

We trained a gLV model on monoculture kinetics and community stationary phase measurements (including three additional passaging timepoints of DTL1 and one additional passaging timepoint of DTL3) to characterize the communities over longer timescales (**Figs. 2b, 3c, S9b-e, Methods**)^17,49,50^. To minimize overfitting of model parameters to the data, we implemented L1 regularization with cross-validation (**Methods, Fig. S10a**). The gLV model was predictive of randomly withheld training data (Pearson rho=0.91, P=3e-83, **Fig. S10b**). In the inferred parameter set, BH positively impacted the growth of ER (**Fig. S10c,d**). This result is consistent with the underprediction of ER by the CSLE model, in which species interact only via competition (**Fig. 2d**). This suggests that the overgrowth of ER in DTL 1 may be a result of the high inoculum densities of ER and BH in comparison to the relatively low inoculum densities of several species (BU, CA, DL and PC) with which ER competes (*a_ER_j_* < −0.25) (**Figs. 3c, S10a**). BH, an acetogen, has been shown to enhance the growth of a similar butyrate producing Firmicute species via metabolite cross feeding^51^. BL received negative interactions from all species excluding PJ, and these interactions summed to the largest negative value among all species (**Fig. S10e)**. This suggested that the persistent low abundance of BL, despite its robust monoculture growth and high inoculum densities, can be attributed to the aggregate effect of many negative interactions in the inter-species interaction network.

We designed low and high temporal variability sub-communities over the timescale of four passages to evaluate whether the gLV model could predict distinct classes of dynamic longer-term behaviors. Communities with low temporal variability in community composition could be useful to reduce the frequency of species takeover and/or extinction during dynamic bioprocess strategies, such as fed-batch or continuous cultures, which are commonly used to improve production efficiency^52–54^. Temporal variability was defined as the sum of the Euclidean distances of the relative species abundance between adjacent passages (Methods, equation 14). Low temporal variability communities were identified by maximizing an objective function of the ratio of the Shannon diversity to the Euclidean distance across passages (Methods, equation 15). We used optimization techniques to maximize this objective function across a wide range of initial conditions for all possible (967 total) 3–9-member sub-communities. Notably, among the 967 optimal solutions, only 33 sets of initial conditions displayed unique endpoint species compositions within a small numerical tolerance (**Fig. S11**). We selected a subset of higher diversity unique solutions for experimental validation that represented all species. To determine if the model could distinguish between low and high temporal variability behaviors, we included three representative communities with predicted high temporal variability (i.e., high Euclidean distances) (**Methods**).

Consistent with the model prediction, communities designed for low temporal variability had significantly lower Euclidean distances between passages than communities designed for high temporal variability (p=8e-6, unpaired t-test) (**Fig. 4b**). In addition, the model accurately predicted several qualitative characteristics of the high temporal variability communities, including the highest abundance species at each endpoint (**Fig. 4c**). The model forecasted that FP is outcompeted (greater than 10-fold lower relative abundance in final passage than the initial passage) in the two high-temporal variability communities containing FP (**Fig. 4c**). Notably, the model also identified a low temporal variability subcommunity (**Fig. 4d**, BH-EL-FP) in which FP persisted at a constant relative abundance over passages two through four. BL persisted at a constant relative abundance across the last three passages when cultured with BU and CA (**Fig. 4d**, BU-BL-CA). By contrast, BL displayed low relative abundance in the first passage and high relative abundance in later passages of the 10-member community training data (**Fig. 3c, S9e**). The model predicted four sub-communities in which at least three species persisted at relatively constant relative abundance for at least three passages (BH-EL-FP, BL-BU-CA, CA-EL-ER-PJ, and BU-CA-EL). This demonstrates the gLV model can be used to design communities that display species coexistence over longer timescales.

**Figure 4.**
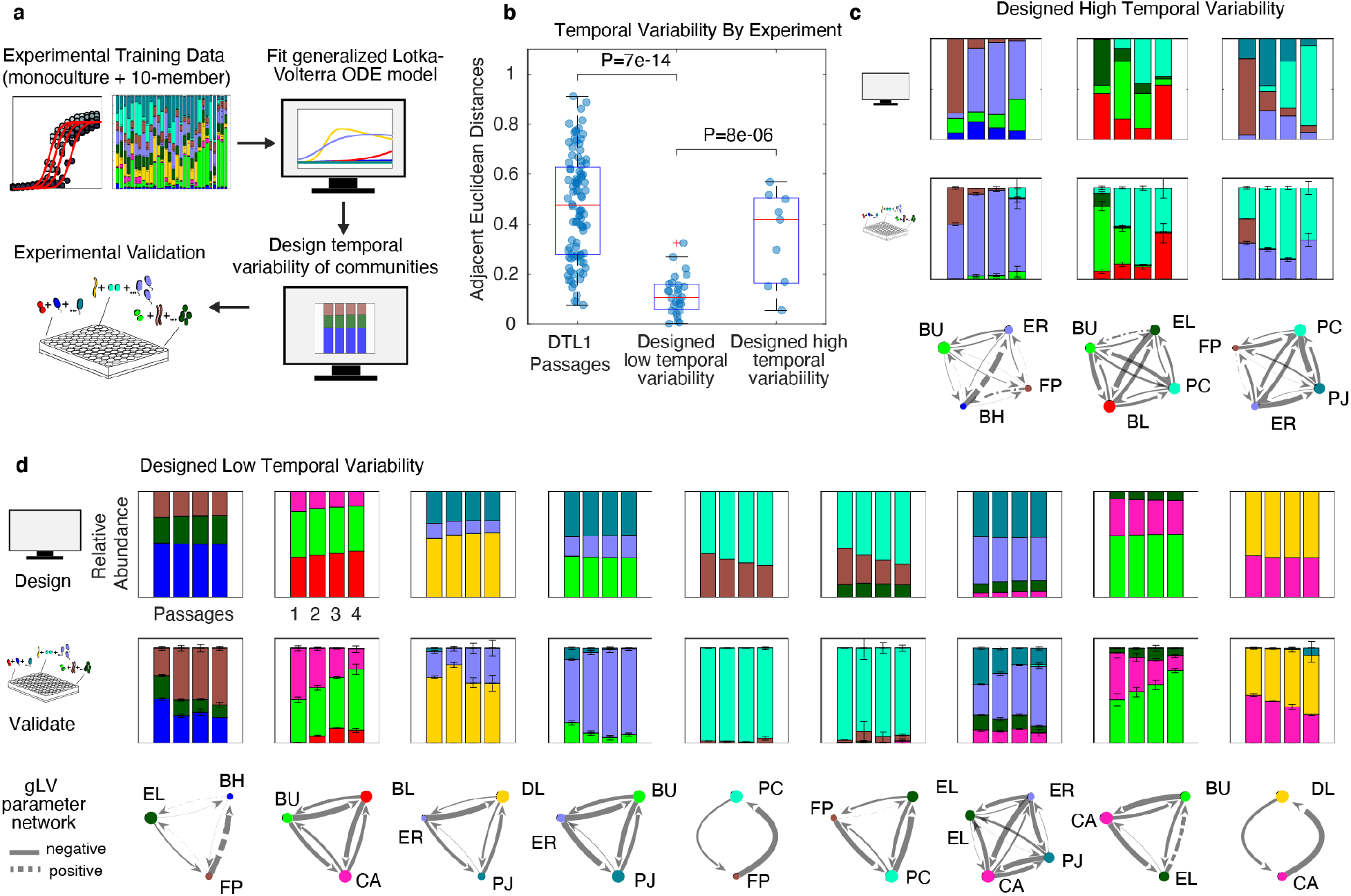
Model-guided design of high and low temporal variability of species composition. **a** Schematic of the experimental workflow where the generalized Lotka-Volterra (gLV, Methods, equation 13) model is trained on monoculture and 10-member community passaging timepoint data (Methods). The gLV model is used to design sub-communities (3-to-9-members) that display low temporal variability of species composition across four simulated passages. Optimization techniques are used to solve for passage 1 initial conditions (i.e., inocula) that maximize the ratio of the summed Shannon diversities to summed Euclidean distances between consecutive stationary phase timepoint measurements (Methods, equation 14, 15). Based on this metric, a set of communities were selected for experimental characterization. **b** Categorial scatter plot of the changes in community composition between passages for DTL1 training data (**Fig. S9a-c**), designed low, and designed high temporal variability communities. Data points denote Euclidean distances between stationary phase community compositions (mean of n=3 biological replicates with outlier detection, Methods) of consecutive passages (Methods, equation 14). Box plot red central line denotes median, upper and lower edges denote 75^th^ and 25^th^ percentiles, respectively, whiskers denote range of non-outlier datapoints, and red “+” denotes outlier. Unpaired, two sample t-test is used to calculate p-value between groups of communities. **c,d** Stacked bar plots of gLV model predictions of stationary phase species composition (top row), experimental measurements (middle row), and inferred gLV inter-species interaction networks (bottom row) for a set of high (**c**) and low (**d**) temporal variability sub-communities. Species color legend follows node labels. Each subplot denotes the relative species abundance at stationary phase of the four passages; for experimental data bar height and error bars denote mean and 1 s.d. of biological replicates (n=3 with outlier detection, Methods). Solid and dashed edges indicate negative and positive inter-species interaction parameters (*a_i,j_*), respectively. Edge width is proportional to the magnitude of the inter-species interaction parameter and node size is proportional to the specific growth rate parameters in the gLV model (*μ_i_*). All biological replicates and omission of 5 cross-contaminated replicates are indicated in Fig S14.

The gLV model trained on monoculture and 10-member community data was moderately predictive of the absolute species abundance in the experimentally characterized 2-4 member communities (Pearson rho=0.56, P=9.6e-10, **Fig. S12**). The unexplained variance in the dataset could be attributed to differences in species richness in the training (10-species) and test data (2-4 species)^25,55^. In sum, these data demonstrate that the gLV model can guide the design of communities that exploit inter-species interactions to support the persistence of lower fitness species over longer timescales, as well as mitigate overgrowth of high fitness species. Therefore, the gLV model informed by variation in inoculum densities of constituent community members of a fixed community size was useful in the prediction and design of community temporal behaviors.

## DISCUSSION

We demonstrate that, despite their complexity, microbial communities are engineerable systems that respond predictably to changes in media formulation and inoculum densities. We develop a data-driven dynamic and statistical modeling framework for tuning these control inputs to optimize the endpoint Shannon diversity of a synthetic human gut community. Using this approach, we increased the Shannon diversity of a representative 10-member synthetic gut community from 53% to 91% of its maximum possible value (**Fig. 3f**). Our DoE and ecological modeling approaches map control inputs to community composition without the need for characterizing detailed biochemical mechanisms (e.g., specific metabolic pathways or metabolites mediating inter-species interactions). As such, the workflow can be generalized to a wide range of communities. Future work could examine effects of monoculture versus community production of live microbial therapeutics on strain engraftment in the host. Since the ecology of the community culture restricts species to their most favorable niches, and can even recapitulate *in vivo* functional profiles, therapeutic communities produced via community culture could be better primed to colonize the competitive gut environment than those produced in monoculture^56–58^.

Our designed media and inoculum conditions yield similar community composition at 500-fold volumetric scale up, suggesting that lab-scale results could translate to production. Though community dynamics are complex, our culture strategy is simple (static, batch culture, no pH control), thus reducing the number of key scale-up parameters. We note that we maintained equivalent headspace gas composition and surface-to-volume ratio during scale up; future studies could confirm whether these are important parameters for anaerobic community scale up. Overall, this efficient, scalable blueprint for designing community assembly should help to alleviate the production bottleneck that limits manufacturing of therapeutic communities at clinical, commercial, and global health scales.

In each stage, we exploit high-throughput, monoculture experiments to first understand the “parts” of our ecosystem, and show that this information is useful for guiding community design. We demonstrate that maximizing monoculture-diversity substantially increases community diversity (**Fig. 1h,i**), and that major trends in community assembly can be explained by constraining monoculture kinetic models with an upper limit on total growth (**Fig. 2**). Model-guided prediction of community assembly from monoculture kinetics allowed us to achieve our design objectives while limiting the number of community measurements. Due to the DNA sequencing pipeline required to analyze species-level composition, community experiments are laborious in comparison to their fully-automated monoculture counterparts. Monoculture-informed prediction of a narrowed initial design space resulted in identification of a high diversity condition within three design-test-learn cycles (**Fig. 3h**).

This ability to rationally inform community experiments with high-throughput monoculture data should make our approach useful for larger communities, potentially even up to 100 members^59^. As species richness increases, the degree of metabolic similarity among species would increase (i.e., metabolic redundancy), leading to potential challenges in identifying specific nutrients that can tune the growth of individual species in the community. However, media design variables could be selected to favor resources such as fibers, peptones, and mucins, which have been shown to support high richness cultures from stool sample inocula^57,60^. In addition, the ability to control the endpoint abundance of each species as a function of its initial density may decrease due to enhanced strength of ecological competition. In this case, our computational modeling and optimization workflow could be modified to identify optimal strategies for partitioning a high richness community into a minimal number of sub-communities that enable control of species via media factors and inocula.

One limitation of our approach was that inoculum density was an insufficient control point for BL, which was subjected to a disproportionate number of strong negative interactions in the community (**Fig 3c**). Despite the robust growth of BL in monoculture (**Fig. 2b**) and a high inoculum density (**Fig. 3b**), this species did not grow well in communities (**Fig. 3c**). To address this limitation, future efforts could use dynamic modeling to leverage multiple inoculation timings as an additional control point for community composition. Design of species-specific inoculation timings would allow for precise manipulation of inter-species interactions over time. In the simplest case, a species that does not grow well in communities due to negative interactions could be given a “head start” by inoculating at an earlier timepoint. Further, a “temporary support community” could be designed to boost the initial growth of a low fitness species prior to inoculating the remaining community members at a later timepoint. Similar approaches could be used to control an organism that tends to overgrow.

As a proof of concept that inter-species interactions can be leveraged to design temporal behaviors, we used a data-driven generalized Lotka-Volterra (gLV) model to guide the design of communities with low variability of species composition over time (**Fig. 4**). We note that our implementation of the gLV model describes batch culture growth (including stationary phase) with non-equilibrium trajectories. Endpoint community composition of batch culture was predicted quite accurately as a function of initial conditions by fitting these transient dynamics to experimental data (**Fig. S9b**). By contrast, theoretical analyses of the gLV model tend to focus on long-term behaviors (e.g. stable steady-states or limit cycles), to which many different initial conditions converge^61^. This nuance between our data-driven implementation and most ecological analyses illustrates that in spite of the constrained long-term behaviors of the gLV model, it is useful for designing specific community compositions as a function of initial conditions.

Defined microbial communities hold significant promise for many applications including agriculture, biofuels, and medicine^62^. We developed a general control strategy for complex microbial communities and applied these strategies to address the challenge of manufacturing defined human gut communities for therapeutic applications. Beyond therapeutic community production, our method will be broadly useful for defined microbial community bioprocesses. For example, in metabolic engineering applications wherein designed pathways are distributed among distinct community members to exploit division-of-labor, our method could be applied to tune community member proportions and thus optimize metabolite product yields^54,63^. Eventually, the ability to identify influential control parameters for steering microbial community composition and functions could be used to modulate an unhealthy patient’s microbiome towards a healthy state. For instance, mirroring media component manipulation, changes in diet are well documented to shape gut microbiome composition. It was also recently shown that dosage strength (i.e. inoculum density) was a critical factor in the successful redesign of the first phase three clinical trial of a donor-derived live microbial therapeutic for treating recurrent *C. difficile* infection^27,64,65^. Overall, initial species densities, environmental resources, and inter-species interactions are key design parameters for engineering microbial community dynamics, from community bioprocessing to potentially designing an ecological restoration of a dysbiotic gut microbiome.

## METHODS

### Strain maintenance, precultures, and growth media

The following methods are adapted Hromada 2021, Clark 2021 and Venturelli 2018^25,28,66^. All anaerobic culturing was carried out in a custom anaerobic chamber (Coy Laboratory Products, Inc) with an atmosphere of 2.5 ± 0.5% H2, 15 ± 1% CO2 and balance N2. All prepared media, stock solutions, and materials were placed in the chamber at least overnight before use to equilibrate with the chamber atmosphere. The strains used in this work were obtained from the sources listed in Supplementary File 1 and permanent stocks of each were stored in 25% glycerol at −80 °C. Batches of single-use glycerol stocks were produced for each strain by first growing a culture from the permanent stock in anaerobic basal broth (ABB) media (HiMedia or Oxoid) to stationary phase, mixing the culture in an equal volume of 50% glycerol, and aliquoting 400 μL into Matrix Tubes (ThermoFisher) for storage at −80 °C. Quality control for each batch of single-use glycerol stocks included (1) plating a sample of the aliquoted mixture onto LB media (Sigma-Aldrich) for incubation at 37 °C in ambient air to detect aerobic contaminants and (2) next-generation DNA sequencing of 16S rDNA isolated from pellets of the aliquoted mixture to verify the identity of the organism (Illumina). For each experiment, precultures of each species were prepared by thawing a single-use glycerol stock and combining the inoculation volume and media listed in Supplementary File 1 to a total volume of 5 mL for stationary incubation at 37 °C. Incubation times are also listed in Supplementary File 1. Prior to inoculating starter cultures, the workspace and pipettes were cleaned with Spor-klenz (STERIS), and again with 70% ethanol between strain inoculations. A clean Kim-wipe (Kimberly-Clark) was held above the workspace to check for air currents from equipment fans that could lead to cross contaminations, and equipment was turned off or rearranged as needed. Anaerobic work was conducted in a spatially linear workflow from cleanest to least clean materials (e.g.) tips, clean reagents, cell containing media, then trash, as ordered from dominant to non-dominant hand. Motions above open, sterile containers is restricted to minimum necessary actions.

### Genomic DNA extraction, DNA library preparation, sequencing, primer design, and data analysis

DNA extraction, library preparation, and sequencing were performed according to methods described in Hromada 2021 and Clark 2021 ^25,66^. In brief, cell pellets from about 150 uL of culture were stored at −80C following experiments. Genomic DNA was extracted using a 96-well plate adaption of the DNeasy protocol (Qiagen). Genomic DNA was normalized to 1 ng/uL in molecular grade water, and stored at −20C. Dual-indexed primers for multiplexed amplicon sequencing of the v3-v4 region of the 16S gene were designed as described previously, and arrayed in 96-well, skirted PCR plates (Thomas Scientific) using an acoustic liquid handling robot (Echo LabCyte). Genomic DNA and PCR master mix were added to primer plates and amplified prior to sequencing on an Illumina MiSeq platform.

Sequencing data were analyzed as described in Hromada 2021. In brief, basespace Sequencing Hub’s FastQ Generation demultiplexed the indices and generated FastQ files. Paired reads were merged using PEAR (Paired-End reAd mergeR) v0.9.0 (Zhang et al, 2014)^67^. Reads were mapped to a reference database of species used in this study, using the mothur v1.40.5, and the Wang method (Wang et al, 2007; Schloss et al, 2009)^68,69^. Relative abundance was calculated by dividing the read counts mapped to each organism by the total reads in the sample. Absolute abundance was calculated by multiplying the relative abundance of an organism by the OD600 of the sample. Samples were excluded from further analysis if > 1% of the reads were assigned to a species not expected to be in the community (indicating contamination).

### Monoculture media screening experiment

The media screening experiment was designed to improve monoculture-diversity (equation 4) on DM38, a chemically defined medium developed in our laboratory, and referenced as the “baseline” medium in the text. Supplementary File 2 contains the medium and stock solution recipes referenced in this section. A four-factor, two-level half factorial screening design with appended center point condition was constructed in JMP 15 (SAS institute). “High” absolute design levels for sugar mixture, amino acid mixture, and pH variables (these are key components in DM38) were set at their respective DM38 concentrations. Yeast extract (sterile filtered, not autoclaved) was included to support monoculture growth of F. prausntitzii, as keenly observed by D’Hoe et al ^41^. “Low” design levels were set at 0 g/L for sugars, amino acids, and yeast extract, and 5.7 for pH (according to generally reported ranges for the human large intestine^70^). Stock solutions of sugars, amino acid mixture, and yeast extract were prepared at 20x v/v of their target “high” concentrations, and sterile filtered. The nine media were arrayed according to the experimental design in 2mL deep-well blocks (Nest), using a Tecan Evo liquid handling robot to aliquot the appropriate volume of 20x stocks into 1.4x base medium. The final concentration was brought to 1x using sterile water. The deep well blocks, containing ten sets of the media experimental design, were inoculated from the ten precultures to a 600nm optical density value of 0.01. Optical density was measured using 200 uL of sample in a Tecan F200 plate reader in standard clear, flat bottom 96-well microplates (Grenier). Inoculation volumes were calculated as Volume_(inoc)_ = Volume_(well)_*0.01 OD / (Preculture OD). Inoculation was performed from a sterile trough with a multichannel pipette. Four 200 uL replicates were mixed and aliquoted to sterile, clear, flat bottomed, 96-well microplates (Grenier), covered with a transparent seal (Breath EZ, Diversified Biotech), and incubated at 37c in the Tecan Evo incubator. Automated OD600 measurements were recorded every two hours for about 60 hours with a Tecan F200 plate reader.

### Modeling monoculture growth

Model-guided optimization of community Shannon diversity (equations 1,2) was performed by modeling monoculture growth response (3) on various media. “Monoculture-diversity” (equations 4,5) was used as a proxy function for Shannon diversity, enabling a monoculture-based approach for manipulating community Shannon diversity.

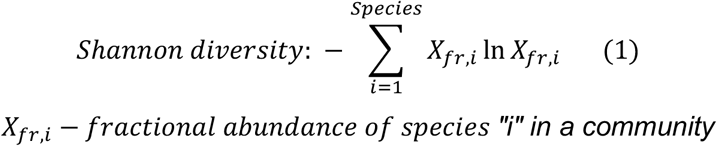

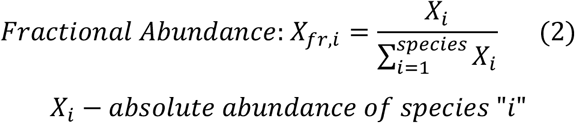

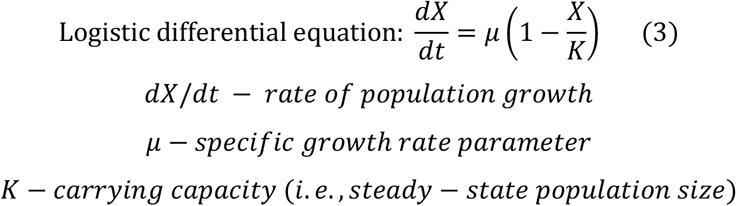

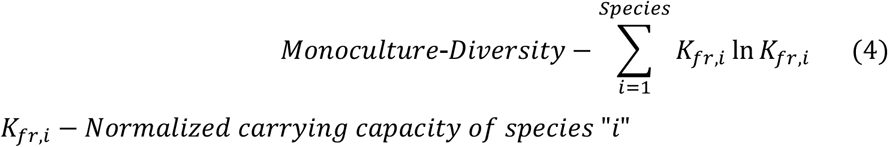

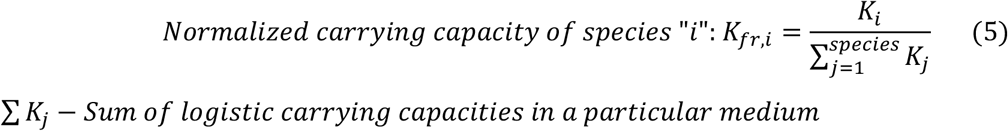

Monoculture timeseries growth data from the media screening experiment was fit with logistic differential equations (equation 3), and the carrying capacity parameter was used as a readout of growth response. Carrying capacity serves as a “smoothed,” time independent maximum growth value. Smoothing is required because raw data may contain outlier values due to condensation on the transparent plate seal or other technical variability. If computational resources or expertise are limited, the growth response could also be taken as the maximum value of a smoothed timeseries (e.g. after applying a running average filter). The baseline of the OD600 timeseries data was computationally “blanked” (i.e. normalized) to the known inoculum density by subtracting the difference between the time-zero measured value and known inoculum from the entire timeseries. Each fitting was performed independently using bounded, nonlinear regression with MATLAB’s “fmincon” function, which returns the logistic parameter set (*μ,K*) that minimizes the sum of squared errors between the model predictions and the experimental data. All timeseries were truncated to 30 hours to remove death phases. Outlier detection was performed by comparing the z-score of the mean OD600 across replicates, to omit replicates that did not grow.

Multivariate polynomial regression models (equation 6) were fit to predict each specie’s carrying capacity parameter (growth response) as a function of the scaled media design matrix (predictors).

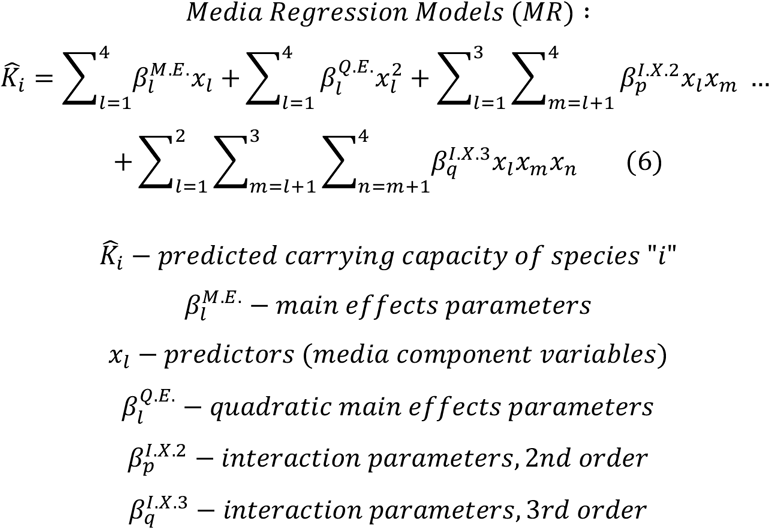

We note that although the model is a multivariate polynomial function of the design variables, the regression is linear with respect to the parameters, as the higher order predictors are treated as “new” variables whose value is calculated prior to regression. The polynomial structure (equation 6) contained main effects (X_1_), quadratic main effects (X_1_^2^), and both second and third order interaction terms (X_1_*X_2_ and X_1_*X_2_*X_3_). The double and triple sum terms in this equation represent the upper triangular matrix of unique two-factor interaction parameters and three-dimensional upper triangular matrix of third order interaction parameters (X_1_*X_2_=X_2_*X_1_ so only one of these predictor terms should be included). The estimation of quadratic terms is contingent on the inclusion of a center point condition in the otherwise two-level experimental design. Because the models are data limited, elastic net regularization and nested cross validation were performed to reduce overfitting. The elastic net and regularization coefficient hyperparameters were selected using a “grid search” approach, and MATLAB’s “lasso” function. For each species, the 9-condition dataset (9×16 predictor matrix and 9×1 growth response vector) was partitioned into all nine possible combinations of eight conditions (rows) using MATLAB’s “crossvalind” function (first partitioning). The “lasso” function is called with the cross-validation argument, wherein it internally performs a second round of leave-one-out cross validation to identify the regularization and elastic net coefficients (hyperparameters) that minimize the out-of-fold mean sum of squared errors for the “internal” cross validation sets. Only the hyperparameters, but not the regression parameters, are returned at this stage. The Lasso function is then called again without the cross-validation arguments, receiving the previously identified hyperparameters as arguments to find a best fit parameter set for the “first partitioning” of the original dataset. This is performed for each partition of the original dataset, such that each regression model is an ensemble model with nine parameter sets, each corresponding to one “leave-one-out” partitioning of the data. Each parameter set has its own, independently identified hyperparameters, such that none of the hyperparameters are biased by training on the entirety of the dataset. The models are validated by making “out-of-fold predictions”, meaning using the parameters trained on each of the nine partitions of eight datapoints to predict the one datapoint that is not contained in that partition. When the models are called to make a new prediction (e.g. for the optimization script), the nine predictions of the “ensemble” are averaged to a scalar value.

### Media optimization

A constrained optimization problem was solved using MATLAB’s “fmincon” function to solve for the concentration profile of sugar mixture, amino acid mixture, yeast extract, and pH that maximized the monoculture-diversity (equations 6, 7, and 8).

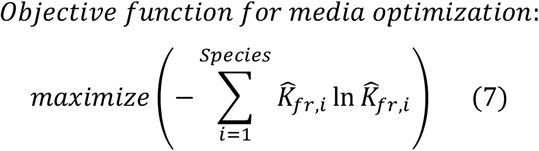

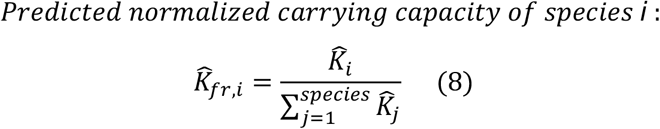

The upper and lower bound arguments to the “fmincon” function are set such to constrain the solution within the original experimental design levels (sugars between 0 and 9.45 g/L, yeast extract between 0 and 2 g/L, amino acids between 0 and 10.7 g/L, and pH between 5.7 and 6.7). The function is initialized with a random guess of the sugars, amino acids, yeast extract, and pH concentrations. The “objective function” references the received concentration inputs and calls the linear regression models to make a prediction of each specie’s carrying capacity from this set of media component concentrations. From these ten carrying capacity predictions, the predicted monoculture-diversity is calculated. The “fmincon” function then iteratively solves for the single concentration of the resources that maximizes the predicted monoculture diversity, using the default interior point algorithm.

### Monoculture growth kinetics over a range of inoculum densities

Deep well blocks (96-well, 2mL, Nest) were filled with 1000uL of the optimized medium. Species were precultured and inoculated into each of the first ten wells of the first row of the block at a density of .01 OD600 as previously described. A multichannel pipet was used to mix and perform six 10-fold volume/volume serial dilutions of the first row down the rows of the plate. Three replicate 96-well microtiter plates with 200uL in each well were aliquoted from the deep well block and covered with a transparent seal, breathable seal. Plates were incubated and timeseries OD600 was recorded as previously described.

Timeseries data from inoculum conditions that did not result in reproducible growth were omitted from the dataset, and data was normalized as previously described. The low inoculum densities resulted in growth curves that “appeared” to have a long lag phase, but were much more likely to be in exponential growth phase at a biomass density that was far below the limit of detection of the plate reader. The exponential and stationary phase data from each specie’s set growth curves was isolated as values greater than the assumed 0.05 lower limit of detection for the plate reader. The true limit of detection of the reader is .001, but data below ~.05 has high signal-to-noise ratios for automated microbial growth. As such, the “measured” initial conditions were omitted from the dataset, as they generally reflected the low limit of detection of the platereader. Nonlinear regression was used to solve for the single logistic parameter set (*μ,K*) and the set of initial conditions (one for each growth curve in the set) that minimized the sum of squared errors between the model predictions and the exponential phase data. A vector of two logistic parameters and one-to-six initial conditions (depending on how many dilutions grew reproducibly) was passed as variables to the “fmincon” solver. The objective function then parsed the vector into initial conditions and ODE parameters, then called an ODE solver to generate model predictions. The value of the objective function is the sum of mean squared errors between the model predictions and the exponential phase data for all growth curves in the set. The “fmincon” function returns the vector of parameters and initial conditions that minimize the objective function. The computationally fitted initial conditions were plotted in log-log space against the experimental initial conditions, and a first order linear regression was performed to map the log transformed experimental initial conditions to the log transformed, computationally fitted initial conditions, using sets of values that fell in the linear range.

### Design of the first community inoculum density experiment (DTL1)

The experimental design chosen for the first inoculum screening was a nine-factor, three-level definitive screening design^71^. These designs have three levels for each variable, improving estimation of the quadratic effects that are likely important for approximating the endpoint of exponential microbial growth with a polynomial function. The scaled design matrix was constructed in JMP 15. Inoculum concentrations were assigned to the scaled experimental design levels using solutions from the constrained system of logistic equations model. The constrained system of logistic equations was simulated in MATLAB, using the growth rate and carrying capacity parameters as fitted to monoculture data (described in the previous section).

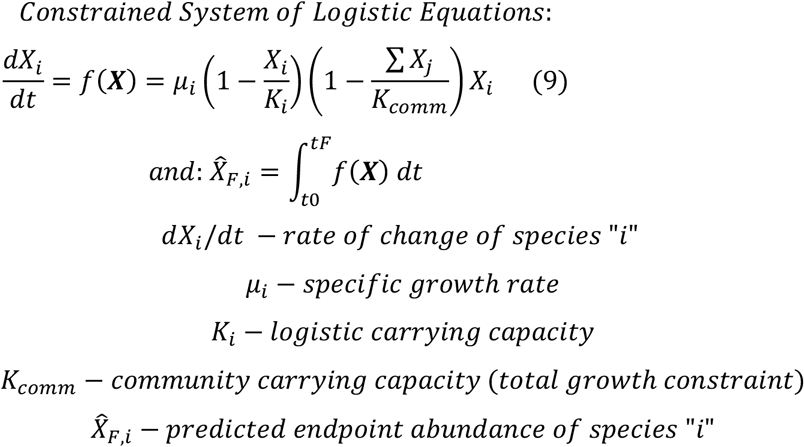

The community carrying capacity parameter *K_comm_* was taken as the maximum OD600 of a full community culture inoculated from an even inoculum (all species inoculated to .001 OD600). To find the set of initial conditions that maximized the Shannon diversity of the CSLE model at steady state, a constrained optimization problem was solved with MATLAB’s “fmincon” function. The variables optimized by the “fmincon” solver consisted of the set of all species’ initial conditions. The objective function internally maps these initial conditions to the computational space equivalent (using the linear regression functions previously described), and simulates community growth by calling a CSLE ODE function. The “fmincon” solver solves for the set of initial conditions that maximize the Shannon diversity (equation 1) of the steady state population abundances using the default interior point algorithm.

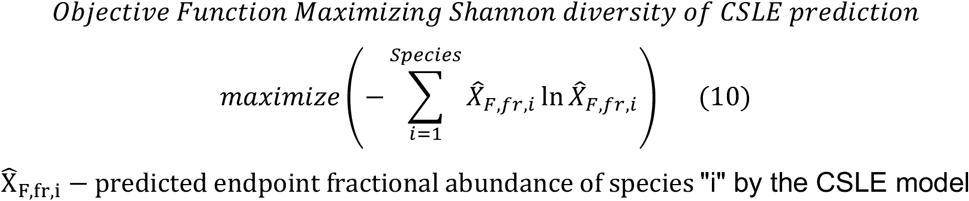

The initial condition solutions are constrained by lower bounds of the experimental inoculum conditions that did not grow, such that the solver does not return initial condition that are too low to use in practice (an issue that can arise when modeling populations as continuous numerical variables). The total inoculum is constrained using a linear inequality argument such that the sum of all initial conditions did not exceed 0.02 (*F. prausnitzii* was fixed at 0.01; the sum of the other nine species was constrained to below 0.01). The high inoculum level for each species was solved for by fixing all other species’ initial conditions at the maximum diversity solution (center point), then finding the initial condition for that species which yielded a 3.3-fold higher steady state abundance than the center point condition. Specifically, “fmincon” was called to minimize the squared error between the simulation and 3.3 times the steady state abundance of that specie’s maximum diversity solution as a function of that species initial condition. This was iteratively performed to find all species’ “high” initial condition levels for the experimental design. The low levels were set symmetrically to the “high” levels in log space, (e.g. the center point was multiplied and divided by the same x-fold factor), such that a CSLE simulation of the experimental design conditions predicted maximum diversity at the center point, and an approximate a 10-fold range of steady state abundances of each species occurred between “high” and “low” design levels. This approach accounts for the fact that a species with a very fast exponential growth rate will likely need a much larger perturbation (in comparison to a species with a low exponential growth rate) to its initial condition to achieve a similar change in the endpoint growth.

### Community inoculum density experiments

Experimental designs were arrayed with a Tecan Evoware liquid handling robot. Before inoculation, precultures were centrifuged at 4000 rpm, 7.5 minutes in a Sorvall ST 16R centrifuge (Thermo Scientific). Anaerobically, the supernatant was decanted, the pellet was dry-vortexed, and resuspended in fresh optimized medium using a serological pipette (Drummond). Two 24-well blocks were used to array various densities of the precultures. The top row contained a high-density preculture, the second row contained a mid-density preculture, and the third row contained a low-density preculture. The concentration of the high-density preculture well for each species was calculated by finding the number of ten-fold dilutions of the measured preculture OD which resulted in the smallest inoculation volume greater than 7 uL. In other words, we calculated the lowest volume that can be accurately pipetted by the robot to inoculate the deep well block to its target “high” experimental level. For example, if species A grew to a preculture OD of .2 and was to be inoculated to a target “high” level of .0001 in a volume of 700 uL, then the high-density preculture well would contain a hundred-fold dilution of the preculture (.002 OD600), such that “high” experimental condition would be inoculated with V = .0001 OD * 700 uL / (.002) = 35 uL. This strategy was implemented because any volume less than 7 uL could not be pipetted accurately, while larger inoculum volumes would quickly accumulate and result in a scenario where the sum of all species’ inoculum volumes exceeds the target culture volume. The “mid” and “low” preculture wells were filled by diluting the “high” preculture well by the same x-fold ratio of the high to center point design levels (and equivalently the ratio between the center point and low levels). Two serial dilutions at this ratio were performed from high to mid, and mid to low preculture wells for each species, such that each specie’s high, center point, and low design levels were inoculated with a constant volume from the high, mid, and low preculture wells, respectively. A 200 uL aliquot of the inoculated deep well block was transferred to a 200 uL microplate, covered with a breathable seal, and incubated in the Tecan F200 plater reader at 37C. Labware and culture conditions were consistent between monospecies and coculture, as it should be noted that differences in labware geometries, particularly surface to volume ratios, can affect anaerobic microbial growth dynamics. Optical density measurements were recorded at 28 +/-1 hour in the platereader. 150 uL of the endpoint culture was transferred to a sterile 1mL deep well block and centrifuged at 2400xg for 10 minutes. The supernatant was removed, and the pellet was stored at −80c. 20 uL of the supernatant was used to measure pH using a spectrophotometric phenol red assay, as described in Clark 2021^25^.

### Design of subsequent inoculum experiments

Linear regression was used to fit polynomial models (equation 11) to predict each specie’s community abundance from the inoculum design matrix, using the nested cross validation approach detailed in the media design methods section.

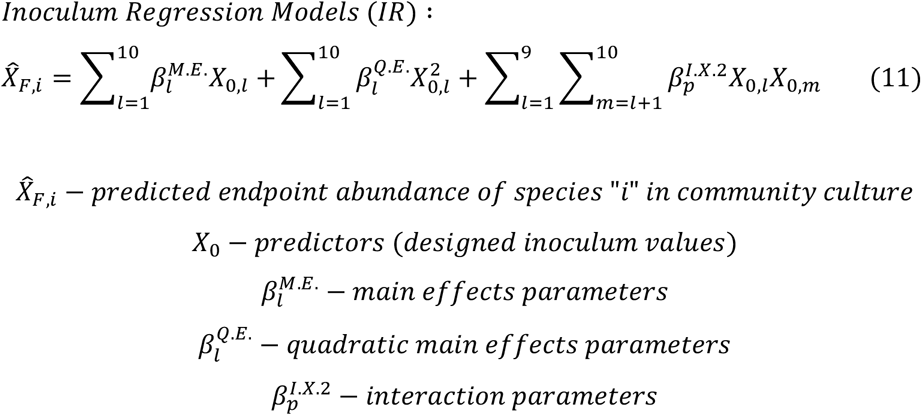

The inoculum design matrix was log10 transformed to scale the values prior to fitting. The models trained on cycles 1+2 community data were evaluated on withheld test data (5 of 59 total conditions) to demonstrate predictivity of the approach (Fig 3B). Replicates were averaged prior to fitting to avoid biasing test/validation data with conditions contained in training data. Validation predictions and Pearson correlation coefficients for both cycles’ models are shown in supplemental materials. Models that were deemed predictive were used in a multi-objective optimization problem (equation 12, details in following paragraph) to predict an updated center point for the new experimental design. Any desired target composition (not only even endpoint, i.e. maximum diversity) can be designed with this approach by updating this target vector with the desired endpoint abundances. Species whose models were not deemed predictive were adjusted using a rational “frameshift” strategy (a graphical representation is provided in supplemental figures). The “frameshift” involves selection of new design level absolute setpoints as follows: if a species overgrew (saturated response) in the previous experiment, the new center point level is set at the previous low level. If a species undergrew (non-measurable or very low growth in comparison to other species), its updated center point inoculum level is set at the previous “high” level. These new center point levels were thus equivalent to the extrema of the previous design space, and could be used as inputs to the regression models (without forcing the models to extrapolate beyond the bounds of training data). We note that the DTL process could probably be carried out using only the “frameshift” strategy to approach a design goal. The magnitude of the levels (x-fold of center point) were maintained between cycles one and two, unless the total range between high and low exceeded two orders of magnitude, in which case it was constrained to two orders of magnitude. In cycle three, the experimental design was modified to a twelve-run Placket-Burman screening with center point, with levels set at two-fold above and below center point. This adjustment of the levels initially informed by the CSLE model (cycle 1 levels) is a qualitative decision that reflects the purpose of the designs. Cycle one had large magnitude levels because it was meant to explore a large design space. Cycle two levels were constrained to two orders of magnitude or less to balance searching the design space with the probability of finding a high diversity condition. Cycle three levels were constrained to only two-fold because the purpose of the design was to demonstrate the robustness of a high confidence prediction to small variations, rather than to explore the design space and gather data for further model training.

A constrained multi-objective optimization problem was solved to minimize the error between target abundances and regression model predictions. This objective function is a more strict definition of maximizing Shannon diversity at a particular total species abundance, and was chosen because maximizing the Shannon diversity can return very low total growth solutions. Additionally, it is also a more flexible approach, as it allows the user to define an exact target community composition. We targeted an even endpoint abundance for each organism of magnitude (average community OD) / (# of species), where the average community OD was the average endpoint OD across all the conditions of the previous experiment.

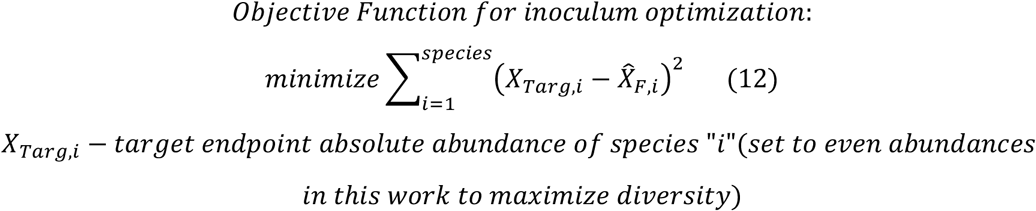

### Passaging experiments

A serial subculture is performed by mixing well and diluting 20 uL of the endpoint community culture into 500 uL of fresh medium (25-fold v/v). The new culture is then aliquoted (200 uL) into a microplate and incubated as previously described. This process was performed three times for the first inoculum design (DLT cycle 1) and once for DTL cycle 2. The data is available in supplemental materials.

### Generalized Lotka-Volterra model training and validation

The parameters of a generalized Lotka-Volterra (gLV) model were fit to monoculture timeseries data and 10-member community initial and stationary phase data.

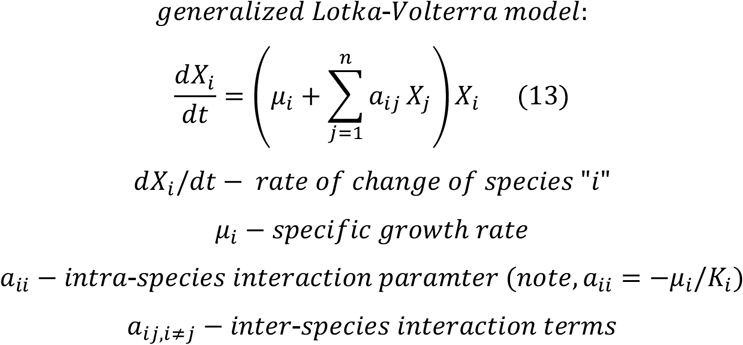

The training data additionally included three passages of the first inoculum screening and one passage of the third, passaging method is described in the previous paragraph. The passages were treated as independent experiments with initial conditions calculated from the previous culture’s endpoint abundances divided by the 25 (corresponding to the volumetric dilution performed to inoculate). The gLV model was fit to experimental data using MATLAB’s “fmincon” solver to minimize a cost function as a function of the model parameter values. The cost function consisted of the sum of squared errors between the model predictions and data, with an L1 regularization penalty to minimize overfitting, as previously described^28^. The upper bounds for growth rate terms *μ_i_*, self interaction terms *a_ii_*, and interspecies interaction terms *a_ij,i≠j_* were 3, 10, and 0, respectively. The lower bounds for these quantities were 0, −10, and −10, respectively. Self-interaction terms must be non-positive and growth rate terms must be non-negative to avoid divergence and maintain biological meaning. The “MaxFunctionEvaluation” and “MaxIterations” arguments for “fmincon” were both set to “Inf” via the “optimoptions” function to allow the solver sufficient time to converge. The solver was initialized with the monoculture growth rates, monoculture derived self-interaction terms, and zeros as respective initial guesses for the gLV growth rates, gLV self-interaction terms, and gLV interspecies interaction terms. Zero is a logical initial guess for unknown parameters subject to L1 regularization, which pushes poorly constrained parameters towards zero. The community data was randomly partitioned into test and training+validation datasets consisting of 10% and 90% of the data, respectively, using MATLAB’s “randsample” function. Monoculture data was not included in validation or test sets because it is collected at high-resolution time intervals, and thus not as strong of a challenge to the model’s predictivity as community data. The regularization coefficient was found by scanning a logarithmic range of values and identifying the value that corresponded to the lowest averaged sum of squared errors across out-of-fold predictions (5-fold cross validation, training +validation data partitioned using MATLAB’s “crossvalind” function). A best-fit parameter set was then re-fitted to the training+validation dataset using the identified regularization coefficient. The model was evaluated for predictivity on the randomly withheld test data. The parameter value heatmap, histogram and, predicted vs. measured scatter plot are shown in supplemental materials Fig S10.

### Design of temporal variability in subcommunities

The best-fit gLV model was used to design communities with low temporal variability over the course of four simulated passages. For all 967 possible 3-to-9-member subcommunities (i.e. sum of 10 choose k for k=3 to 9), a constrained optimization problem was solved to minimize an objective function as a function of the initial conditions of the species present in the subcommunity.

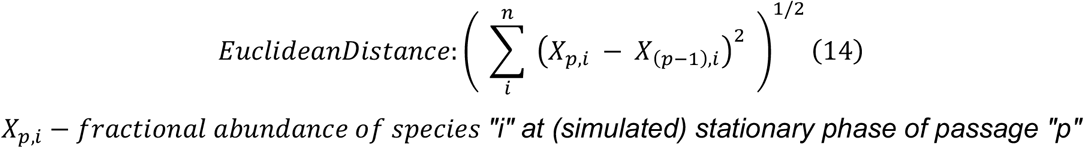

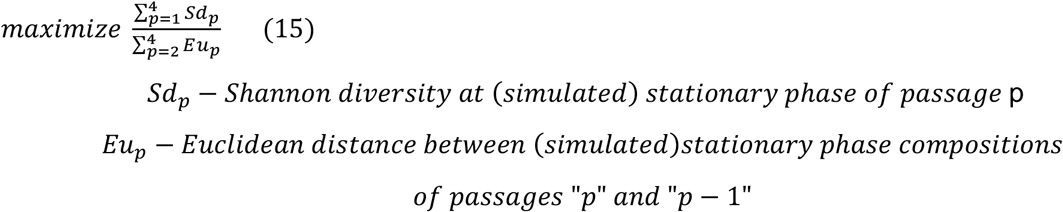

Species absence in subcommunities were simulated by forcing both upper and lower bounds of the omitted species’ population sizes to zero. Initial condition solutions were bounded between zero and 0.01 simulated OD600 for species present in a subcommunity. The endpoint compositions of resulting from all initial condition solutions were sorted into unique results using MATLAB’s “uniquetol” function (within a numerical tolerance of 0.05 for each species). Nine of these communities were chosen for experimental validation on the qualitative criteria of having all species present in the set of subcommunities. These nine communities were of size 2-4 members. As a comparison, we designed four-member high temporal variability subcommunities by maximizing the product, rather than the ratio, of the diversity and distance terms in equation 15. The nine low temporal variability subcommunities and three high temporal subcommunities were inoculated at densities according to the computational predictions. These inoculum conditions spanned orders of magnitude with no symmetry between conditions. The following strategy was used to inoculate these conditions: an “inoculum” 96-well 2mL deep well block was prepared in which each species’ preculture material was diluted to 0.1 in row one. Tenfold serial dilutions were then performed such that preculture material was available for pipetting at a range of .1 to 10^-5^ OD600. The liquid handling robot was assigned to aspirate from whichever well would result in the smallest aspiration volume greater than 7 uL, for each species in each condition. The culture was incubated, passaged, and sampled as previously described.

### Data Exclusion

The following replicates were omitted from NGS analysis due to cross-contamination of >1% of total reads and/or low total sequencing reads <10% of average: **Fig S14d.i** passage 2 replicate 1 and passage 4 replicate 3, **Fig S14d.vi** replicate 2 passages 2-4. The following growth curve replicates were omitted from logistic analysis in **Fig. 1b** due to lack of growth or suspected contamination, using a z-score threshold of 1.5: BH M5 r1, BH M8 r4, BL M9 r4, BU M3 r1, CA M1 r1, EL M1 r1, ER M1 r1, ER M2 r1, ER M3 r1, FP M3 r4, FP M6 r1, and PC M8 r4. In total, 12 of the 360 replicates across 10 species, 9 media, and 4 replicates were omitted, no more than one replicate was omitted per species/media condition.

## Data Availability

The processed sequencing data and raw optical density data for all experiments are deposited in a Github Repository (https://github.com/bryceconnors/DesignOfCommunityDiversity), which will be made public upon publication. Raw DNA sequencing data will be made available via Zenodo prior to publication.

## Code Availability

Code will be available from GitHub upon publication (https://github.com/bryceconnors/DesignOfCommunityDiversity). Data analysis scripts utilize MATLAB R2020a. Python 3 is used for processing sequencing data. In brief, the data and analyses are organized into sub-folders corresponding to each experiment, each of which contains a ReadMe file. Analysis scripts are contained in “modeling” sub-folders, and load raw or processed data from the “rawData” sub-folders. The “02_ReadMe” file contains instructions for navigating to sections of scripts that produce the plots in the figure panels.

## Acknowledgements

We would like to thank Susan E. Hromada and Yili Qian for helpful discussions. We thank Justin Anderson and Daniel Blasiole of the Wisconsin Alumni Research Foundation for their work in filing the provisional patent application based on this research. Research was sponsored by the Bill and Melinda Gates Foundation and was accomplished under grant number OPP1211893 and National Institutes of Health under grant number R35GM124774.

## Author contributions

B.M.C., O.S.V. and R.L.C. conceived the study. B.M.C carried out the experiments. B.M.C. implemented computational modeling. J.T. assisted with model development. B.M.C., O.S.V. and B.F.P. analyzed data. B.M.C. and O.S.V. wrote the paper and all authors provided feedback on the manuscript. S.J.E. and R.L.C. assisted in experimental data collection. O.S.V. and B.F.P. secured funding.

