## Supplementary Information for "Model-guided design of the diversity of a synthetic human gut community"

#### Table of Contents

**Supplemental Text: CSLE Derivation**……………………..……………………………..……….2-4
**Table S1** – Summary of models referenced in the manuscript………………………………...……5
**Figure S1** - Timeseries measurements and logistic fits for media optimization experiment……..6
**Figure S2** – Interpretation of media regression model parameters………………………………..…7
**Figure S3** - Carrying capacity regression model predictions and statistics………………………..8
**Figure S4** – A note on cross validation for small, sparse experimental designs…………………..…..9
**Figure S5** - Monoculture growth curves from high to very low inoculum densities………....10-11
**Figure S6** - Motivating the fitting of initial conditions for the inoculum density experiment……..12
**Figure S7** - Out-of-fold validation predictions for regression models……………………..……....13
**Figure S8** - “Frameshift” approach to assignment of new inoculum setpoints…………………..….14
**Figure S9** - Passaging data for gLV model training……………………………………………………..…15
**Figure S10** - Hyperparameter selection, predictions, and parameters for gLV model………16-17
**Figure S11** - All unique solutions to low temporal variability optimization……………………..……...18
**Figure S12** - Predictivity of gLV model on designed temporal variability communities……..…..19
**Figure S13** - Stacked bar plots showing biological replicates for DTL cycle and passages……20
**Figure S14** - Stacked bar plots showing biological replicates for benchmark conditions and communities with designed dynamic behaviors……………………………………………………..21

### 1 Supplemental Text: Derivation of the CSLE Model from Mass-action kinetics

We aim to predict the dynamics of community growth given an understanding of monoculture growth kinetics, and an empirical community total growth threshold.

Consider a closed system of bacteria and resources. Let us designate two categories of resource: niche resources and a global resource. Each niche resource is uniquely accessed by a single organism. Niche resources are likely to represent resources used for energy generation, for example arginine consumption by E. lenta. Alternately, a global resource is consumed by all species in a community, at an equivalent yield. This resource designation is likely to reflect nitrogen, phosphorous, trace minerals, etc, which are used by all species to construct biomass, a "substance" who's formulation is quite consistent.

$$X_1 + Y_1^{X/R} R_{N,1} + Y^{X/G} R_G \xrightarrow{\mu_1} 2X_1$$
$$X_2 + Y_2^{X/R} R_{N,2} + Y^{X/G} R_G \xrightarrow{\mu_2} 2X_2$$

And in the $n$ species, $m$ resource scenario:

$$X_i + Y_i^{X/R} R_{N,i} + Y^{X/G} R_G \xrightarrow{\mu_i} 2X_i$$

For $i = 1, 2, ...n$

$X_i$ - species i
$R_{N,i}$ - niche resource i
$R_G$ - global resource
$\mu_i$ - growth rate of species i on niche resource i
$Y_{X/R,i}$ - growth yield of species i on niche resource i
$Y_{X/G}$ - growth yield of all species the global resource

The species' growth rates will then be multiplicative functions of each resource (as in mass-action and multiplicative Monod models). We note that, as in Monod, MacArthur, and Lotka-Volterra models, the yield coefficients are not considered to be true stoichiometric coefficients, and the rate is first order with respect to each resource. In other words, these terms serve to close a population-level mass-balance but do not dictate the molecular-level stoichiometry of the reaction or rate equation.

$$\frac{dX_1}{dt} = \mu_1 R_1^N R^G X_1$$
$$\frac{dX_2}{dt} = \mu_2 R_2^N R^G X_2$$

...for $n$ species and $m$ resources:

$$\frac{dX_i}{dt} = \mu_i R_i^N R^G X_i$$

Niche Resources:

$$\frac{dR_i^N}{dt} = -\frac{1}{Y_i^{X/R}} \frac{dX_i}{dt}$$

Global Resource:

$$\frac{dR^G}{dt} = -\frac{1}{Y^{X/G}} \sum_{i=1}^{n} \left( \frac{dX_i}{dt} \right)$$

Mass Balance on Niche Resource 1:

$$R_1^N(t) = R_{1,0}^N - \frac{1}{Y_1^{X/R}} (X_1(t) - X_{1,0})$$

$X_{i,0}$ - species i initial condition
$R_{i,0}^N$ - niche resource i initial condition

Assuming $X_i(t) >> X_{i,0}$ for simplicity (inocula are usually much lower than total overall growth):

$$R_i^N(t) = R_{i,0}^N - \frac{1}{Y_i^{X/N}} X_i(t)$$

Mass Balance on Global Resource:

$$R^G = R_0^G - \frac{1}{Y^{X/G}} \sum_{i=1}^{n} X_i$$

The yield term $Y_{X/R}$ the amount of biomass a species can produce per unit of resource $R$. In Monoculture, according to the Monod model:

$$Y^{X/R} = \frac{\Delta X}{\Delta R}$$

For batch culture where all resource is consumed:

$$Y^{X/R} = \frac{X_f - X_0}{R_0}$$

For a batch community culture where all species have a constant yield on a common (global) resource), all of which is consumed, we can say:

$$Y^{X/G} = \frac{\sum f_j \Delta X_j}{\Delta R^G} = \frac{\Delta X_{comm.}}{R_0^G}$$

Where $f_j$ is the fraction of the the resource that has been allocated to species j. The sum of all fractions is 1, and $\Delta X$ is the total biomass of any species type that can be produced from an initial quantity of resource, $R_0^G$.

Substituting this expression into the mass balance for $R^G$:

$$R^G(t) = R_0^G \left(1 - \frac{\sum_{j=1}^{n} X_j(t)}{\Delta X_{comm.}}\right)$$

Substituting the above mass balances into the species' ODE:

$$\frac{dX_i}{dt} = \mu_i R_i^N(t) R^G(t) X_i(t)$$

$$= \mu_i \left(R_{i,0}^N - \frac{1}{Y_i^{X/R}} X_i(t)\right) \left(R_0^G \left(1 - \frac{\sum_{i=j}^{n} X_j(t)}{\Delta X_{comm.}}\right)\right) X_i(t)$$

$$= \mu_i R_{i,0}^N R_0^G \left(1 - \frac{X_i(t)}{Y_i^{X/R} R_{i,0}^N}\right) \left(1 - \frac{\sum_{i=j}^{n} X_j(t)}{\Delta X_{comm.}}\right) X_i(t)$$

**Table S1 – Summary of models referenced in main text**

| Model Name | Short Identifier | Independent variable, experimental conditions | Dependent Variable(s) | Terms/Parameters | Description |
|---|---|---|---|---|---|
| Logistic models (media screening) | LM | Single inoculum density, multiple media types | Monoculture growth over time | $\mu$ – specific exponential growth rate; $K$ – carrying capacity | Ordinary differential equation, single population growth model. Fit to timeseries growth measurements for each of 10 monocultures on each of nine media conditions. |
| Media regression models | MR | Concentrations of sugars, amino acids, and yeast extract, and pH | Monoculture carrying capacity | Linear and quadratic main effects, 2nd and 3rd order interactions | Linear regression models predicting each specie's logistic carrying capacity as a function of media component variables. |
| Logistic models (inoculum density experiment) | LI | Range of inoculum densities in optimized media | Monoculture growth over time | $\mu$ – specific exponential growth rate; $K$ – carrying capacity | Fit to each specie's set of timeseries growth measurements for a range of inoculum densities. |
| Constrained system of logistic equations | CSLE | Species inoculum density in a simulated community | Simulated community growth over time | $\mu_i$ – monoculture growth rates; $K_i$ monoculture carrying capacities; $K_{comm}$ – community carrying capacity | System of coupled ordinary differential equations. Predicts community assembly from monoculture as logistic models constrained by a total growth limit. |
| Inoculum regression models (DTL cycle) | IR1<br>IR2 | Inoculum densities from DTL cycles 1, and cycles 1+2, respectively | Endpoint species abundance in community culture | Linear and quadratic main effects, 2nd order interactions | Linear regression models with higher order terms that predict community growth from experimental design of inoculum conditions. |
| Generalized Lotka-Volterra model | gLV | Species inoculum densities in a community culture | Community growth over time | $\mu_i$ – growth rates; $a_{ii}$ – intra-species interactions (note, $a_{ii} = -\mu/K$); $a_{ij}$ – inter-species interaction terms | System of ordinary differential equations describing the growth dynamics of multiple, interacting populations. Trained on monoculture kinetic data from inoculum density experiment (Fig 2), community endpoint data from DTL cycles (Fig3), and endpoint of several passages of DTL cycles (supplementary S8). |

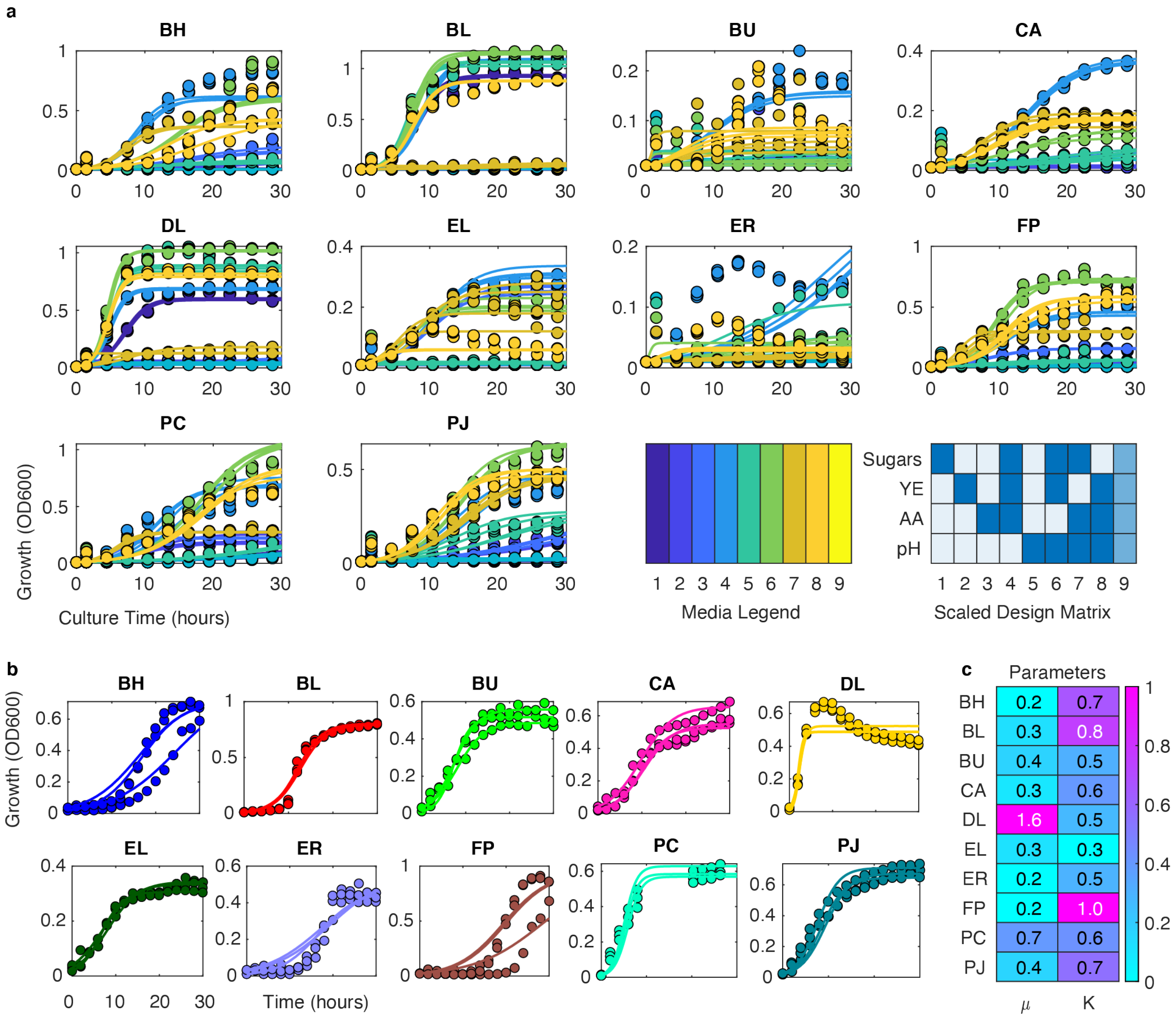


**Figure S1 Timeseries measurements and logistic model fits for media screening experiment and optimized medium.** **a** Scatter plot of timeseries growth (colored circles) measurements for each monoculture (n=3-4 replicates). Color denotes media condition according to figure legend. Lines denote logistic model fits. The absolute concentrations of the components are labeled on the heatmap in Figure 1 of the results section. All species were inoculated to 0.01 OD600. **b** Growth curves and logistic model fits on optimized medium. All species were inoculated at 0.01 OD600. **c** Heatmap of inferred logistic differential equation parameters (mean of 3 replicate fits) for all monospecies in the optimized media: Growth rate ($\mu, hr^{-1}$) and carrying capacity ($K, OD600$).

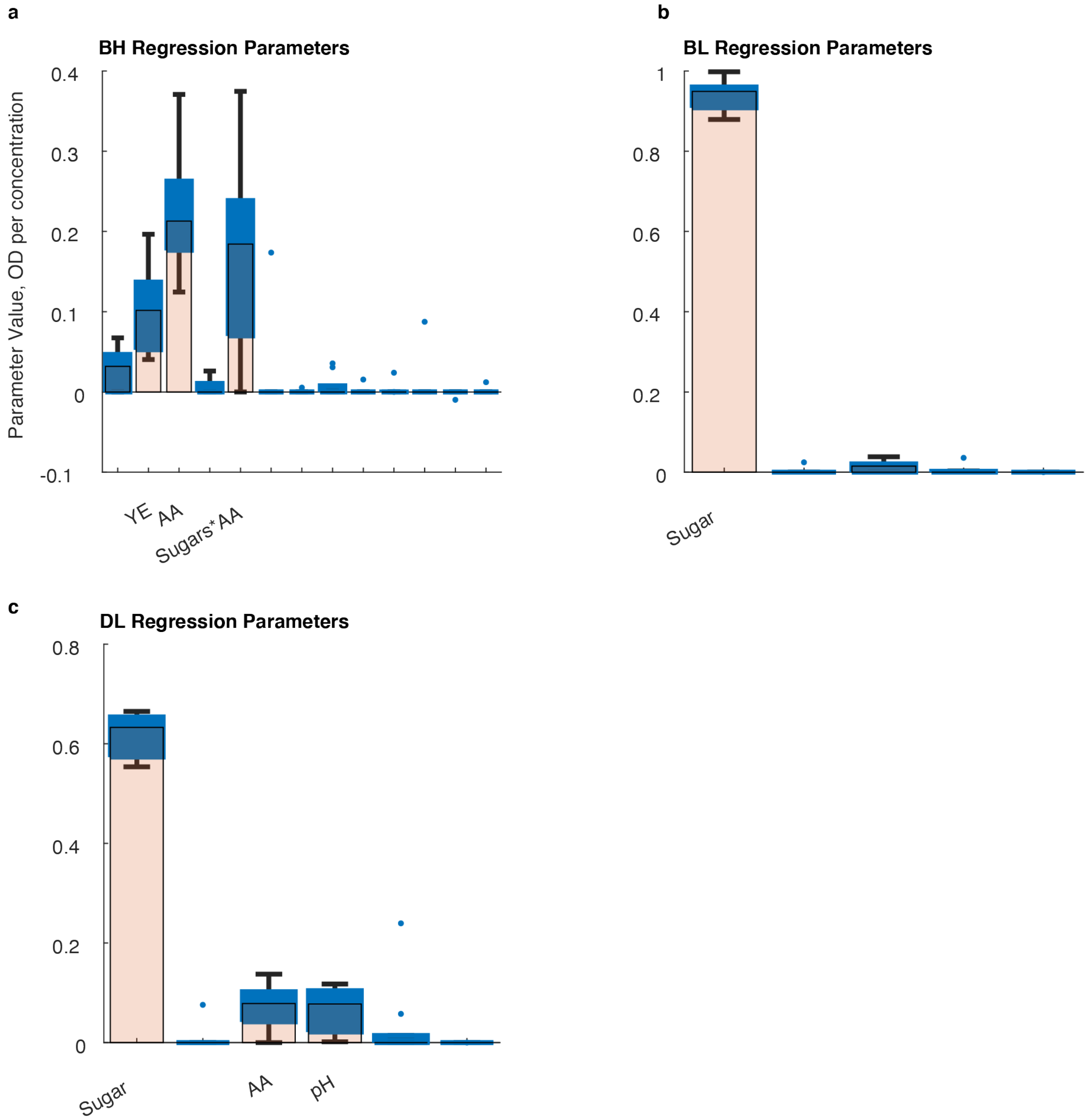


**Figure S2 Media regression model parameters referenced in text**. Boxplots indicate distribution of parameter values for the nine leave-one-out parameter sets for media regression models (MR, Table S1) for **a** BH**, b** BL**, and c** DL. Bar height indicates median value, box upper and lower edges indicate 75$^{th}$ and 25$^{th}$ percentiles, whiskers represent range. Parameters with median values larger than 0.05 have the corresponding predictor labeled on the x-axis. Parameters with a value of zero across all sets are not plotted. All models result from linear regression with linear and quadratic main effects terms, and 2$^{nd}$ and 3$^{rd}$ order interaction terms. Many parameter values are driven to zero by L1 (lasso) regularization. The value on the y-axis represents the effect of each independent variable on the absolute abundance of the indicated species. For example, the BL model (b) effectively consists of only a sugar parameter. Thus, the expected carrying capacity is $\widehat{K}_{BL} = \beta_{sugar} * X_{sugar}$, where $\widehat{K}_{BL}$ is predicted carrying capacity (OD600), $\beta_{sugar}$ is the plotted regression (OD600 per grams per liter), and $X_{sugar}$ is the concentration of sugar in grams per liter.

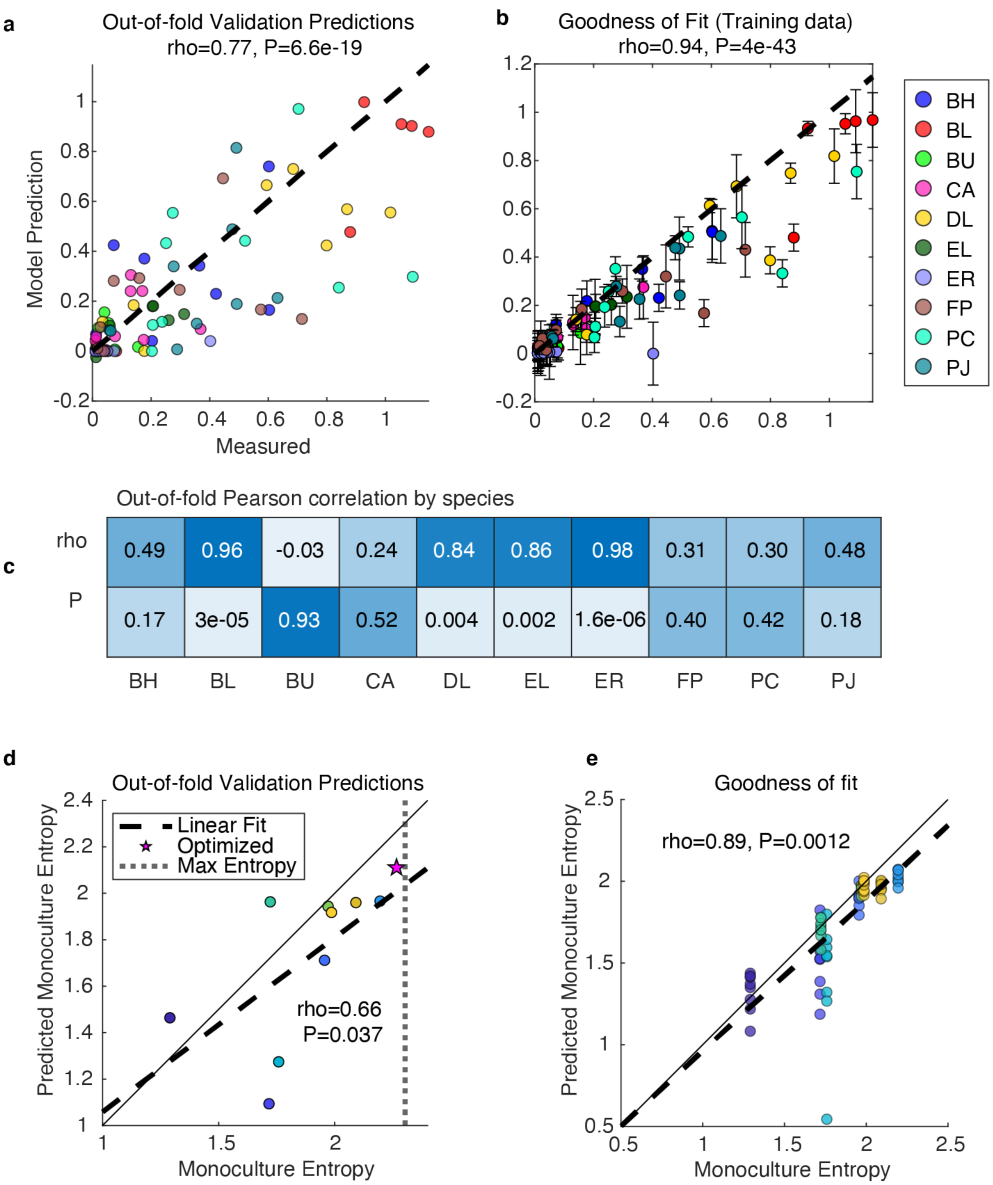


**Figure S3 Media regression model predictivity and goodness of fit.** For each species, a linear regression model (MR, Table S1) is used to predict the individual species carrying capacities for the logistic fits (LM) shown in **Fig. S1** as a function of the four media component variables shown in the heatmap of **Fig. S1**. **a** Scatter plot of out-of-fold predictions for the ten species' media regression models. Nested cross validation is used to fit nine "leave-one-out" parameter sets (leaving out each media condition using cross validation). Pearson correlation (rho) and p-value (P) for all out-of-fold predictions. **b** Scatter plot of model predictions on training data (in-fold predictions). Colored circles represent mean prediction across eight in-fold predictions (i.e., predictions on training data), error bars indicate 1 s.d. from the mean. Goodness of fit statistics (Pearson rho, P) are calculated between the measured data and the mean of predictions on training data. **c** Heatmap of Pearson correlations for out-of-fold model predictions for each species. **d** Scatter plot of predicted vs. actual monoculture-diversity. Predicted values are calculated using carrying capacity predictions on media regression training data. Each of the nine "leave-one-out" parameter sets are used to predict carrying capacity for each of the eight media conditions that are in the training set). Goodness of fit is calculated between the average of the predictions on training data (e.g. across 8 points of a similar color). Pearson correlation coefficient (rho) and p-value (P) are indicated.

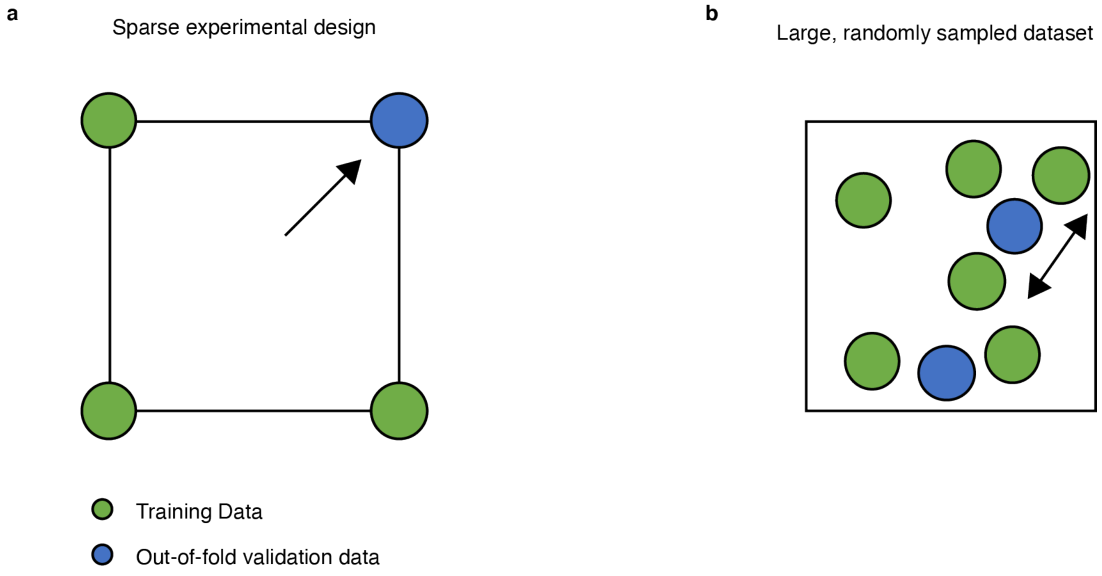


**Figure S4** – **Graphical illustration of cross-validation partitioning for a sparse experimental design dataset vs. a large, randomly sampled dataset. a** Green and blue circles represent hypothetical leave-one-out partitioning for a two, factor, two level design; each node represents a particular condition. To predict the “out-of-fold” point, the model must “extrapolate” (one sided arrow) beyond the range of training data. **b** Hypothetical partitioning of a large, randomly sampled dataset (e.g. a typical machine learning dataset). The model is mostly able to “interpolate” (two-sided arrow) between training data to make predictions on withheld data.

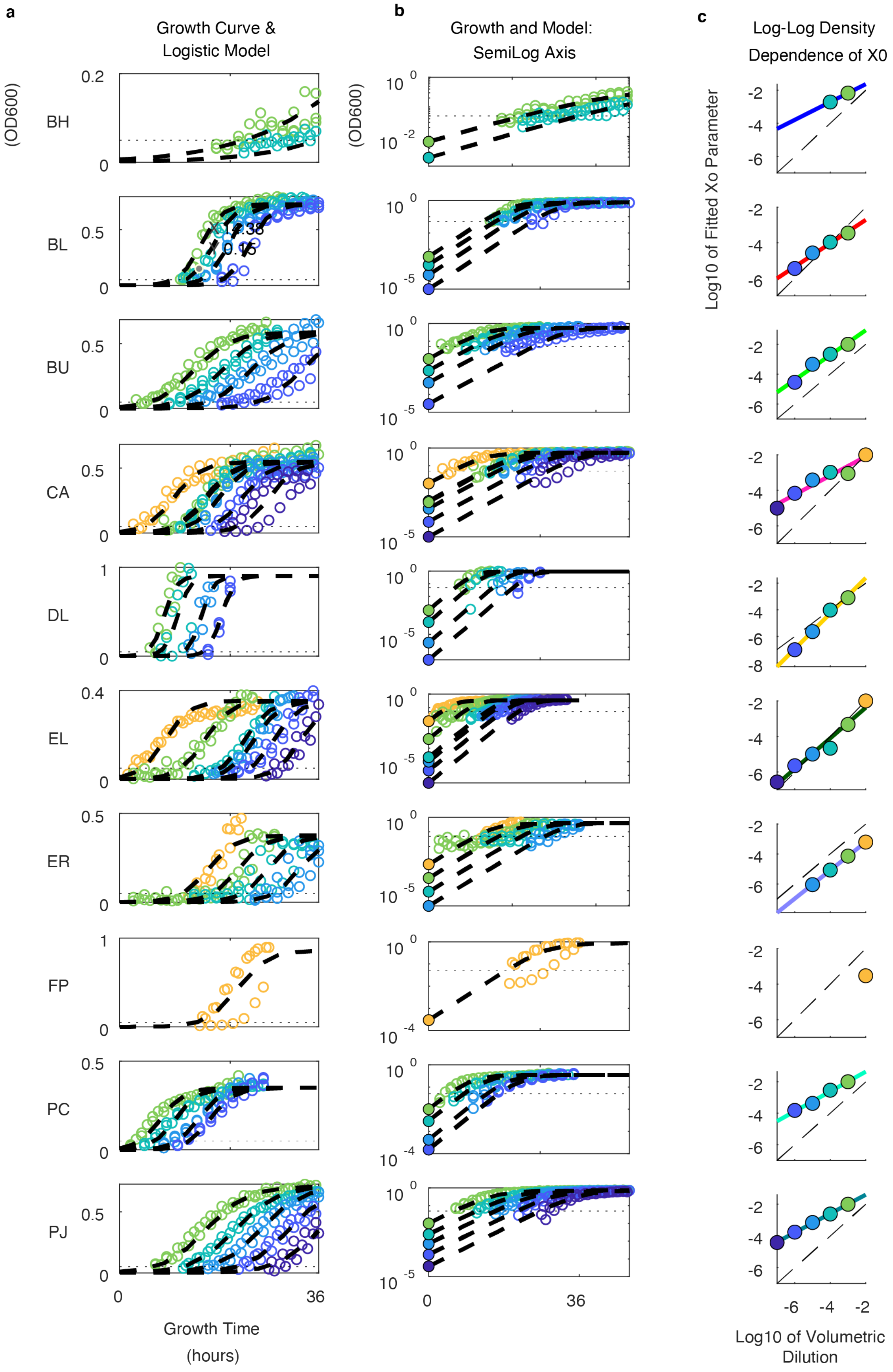

a
Growth Curve & Logistic Model
b
Growth and Model: SemiLog Axis
c
Log-Log Density Dependence of X0
(OD600)
BH
BL
BU
CA
DL
EL
ER
FP
PC
PJ
Growth Time (hours)
Log10 of Fitted Xo Parameter
Log10 of Volumetric Dilution

**Figure S5 Monoculture growth kinetics across a wide range of inoculum densities. a** Scatter plot indicates growth curves resulting from standard inoculum densities (0.01 OD600, yellow) to very low (10E-7 OD600, purple) via 10-fold vol/vol serial dilution. Colored circles indicate 3-4 biological replicates of each monoculture's growth, dashed lines indicate logistic model fits to experimental data (Methods). Dotted horizontal line depicts the approximate low limit of detection (LOD) of the plate reader (0.05 OD600, below this value signal-to-noise ratio is high). **b** The data from (a) are visualized on a semi log axes, revealing that, for lower inoculum densities, the majority of cell doublings are predicted to occur below the low (LOD) of the plate reader (dotted horizontal line). As such, the logistic equation fitting is performed only on data of greater value than the LOD. Since initial conditions are well below the LOD, they are omitted from the training data, and are fitted (i.e. treated as an unknown parameter, similar to growth rate and carrying capacity, Methods). **c** The fitted initial conditions are mapped to the target initial conditions in log-log space, and the inoculum levels that fall on a linear trendline in log-log space are chosen as training data for the CSLE model. This mapping accounts for the following: exponential growth below low LOD, density dependent growth effects, lag phase, and growth which may occur during experimental setup. Though we provide evidence that this method has better goodness of fit (Fig. S6), we note that the increase in accuracy may not merit the complication. A simpler, alternative approach would be to omit timeseries data of value less than 0.05 from training set, append the target initial condition back into the training data, and directly fit the CSLE model to the entire set of all species' monoculture timeseries data (rather than fitting logistic parameters one-at-a-time).

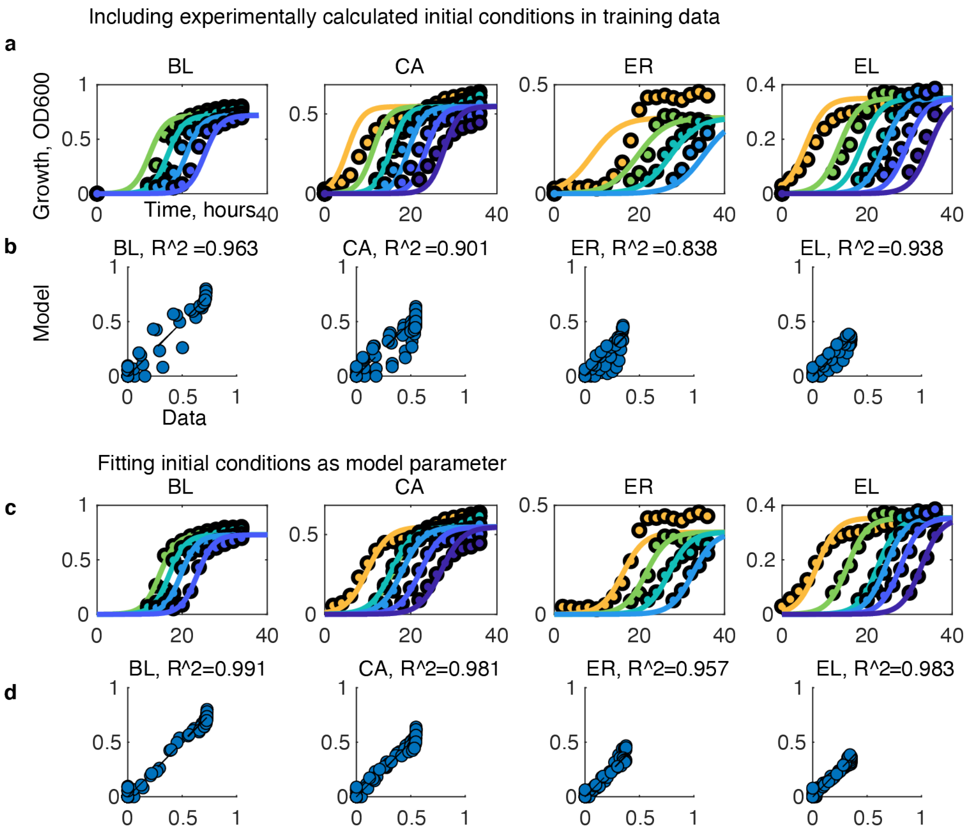


**Figure S6 Motivating the fitting of initial conditions for low the inoculum density experiment**.
**a** Scatter plot of timeseries measurements of OD600 (colored circles) and logistic model fits (lines) for representative species without fitting initial conditions. Yellow represents highest inoculum (0.01 OD), purple represents lowest inoculum (1e-07 OD600), intermediate colors are 10-fold serial dilutions in between. **b** Accuracy of logistic models predictions (LI, Table S1) without fitting the initial conditions. **c** Scatter plot of timeseries growth (colored circles) and logistic model fits (lines) for sample species with initial conditions fitted as a model parameter. **d** Accuracy of predictions for logistic models with initial conditions fitted as a model parameter. The fitting of initial conditions may account for: exponential growth below plate reader limit of detection (LOD), lag phase, and growth which may occur during experimental setup between measuring preculture OD and recording the time zero reading. However, given what we learned throughout the study, we suggest that that the accuracy improvement achieved by this procedure may not merit the complexity, and would suggest simply fitting the constrained system of logistic equations to training data, see S5 caption.

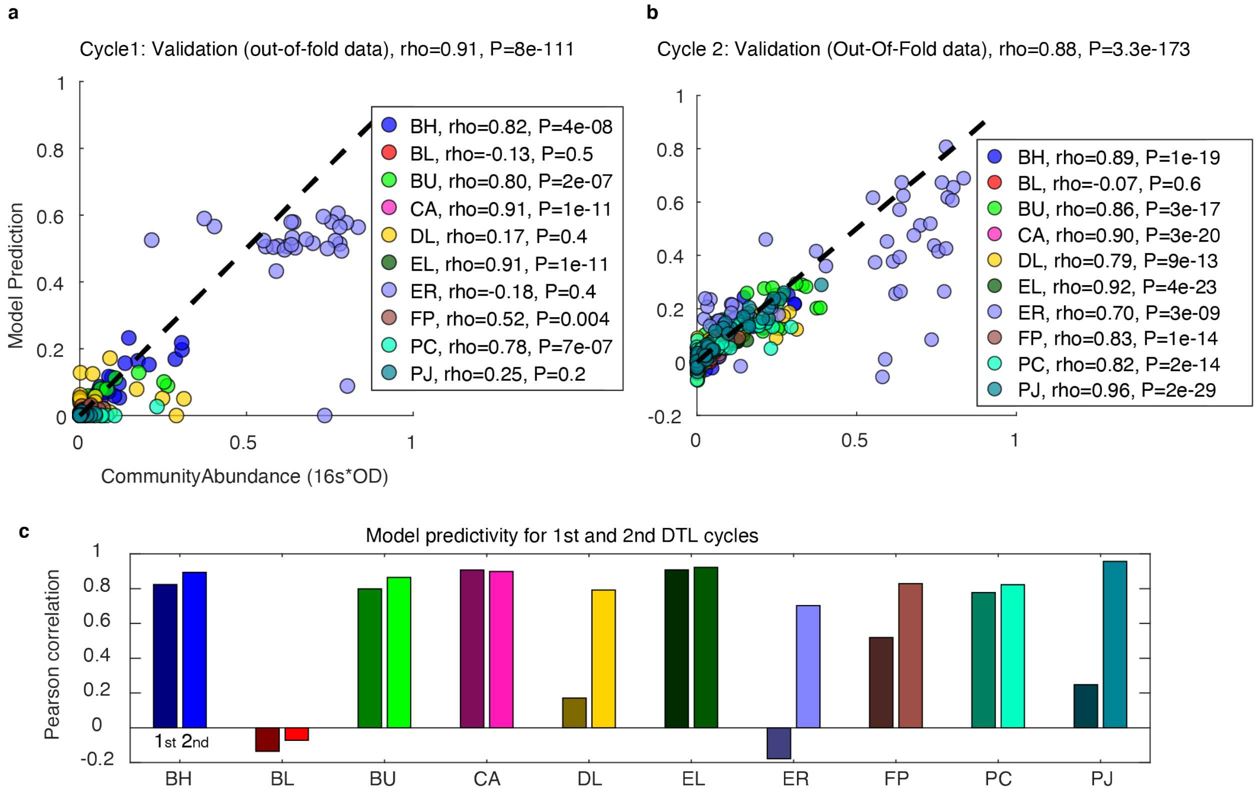


**Figure S7 Regression models for inoculum tuning throughout DTL cycle. a** Scatter plot of inoculum regression model predictions (IR1, models trained on DTL cycle 1 data only, Table S1) vs. experimental measurements (mean of 3 biological replicates) for out-of-fold (validation) conditions/predictions. Pearson correlation (rho) and p-value (P) for each species are indicated in the legend. **b** Scatter plot of inoculum regression model predictions (IR2, models trained on DTL cycle 1+2 data, Table S1) vs. experimental measurements (mean of three biological replicates) for out-of-fold (validation) predictions. Pearson correlations and p-values for each species are indicated in the legend. **c** Bar plot denotes Pearson correlation coefficients after adding DTL 2 training data (IR1 vs. IR2, Table S1). Darker left-hand bars indicate DTL 1 Pearson correlation coefficients (numerical values shown in panel "a" legend). Lighter, right-hand bars indicate DTL 2 Pearson correlation coefficients ("b" legend).

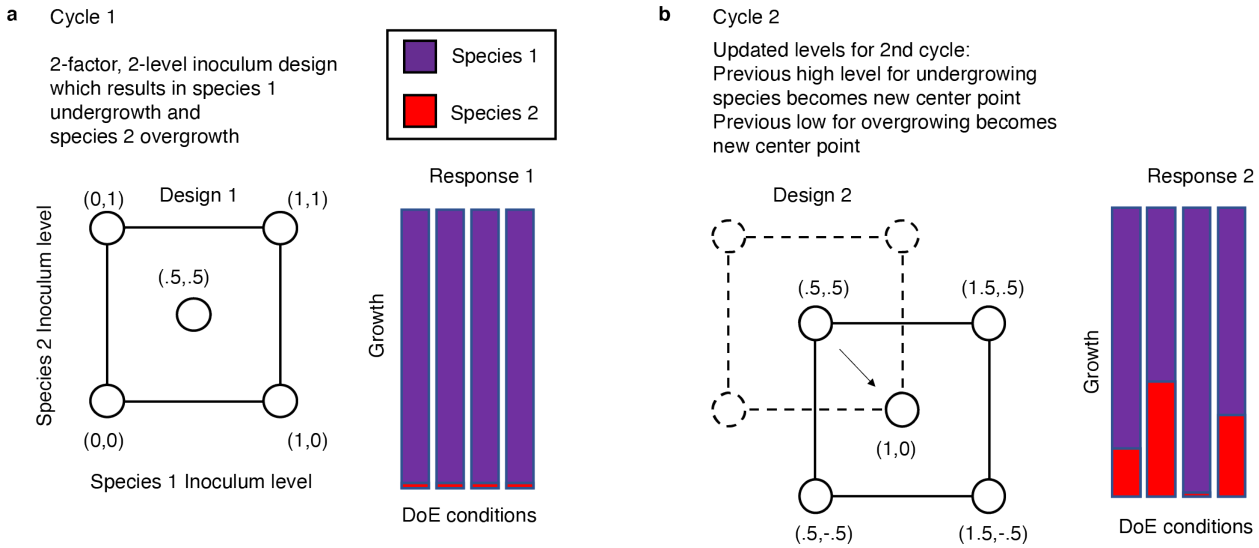


**Figure S8 Cartoon of "frameshift" approach used to assign inoculum values to new experimental designs (used in the case that inoculum regression models were not sufficiently predictive) a** Hypothetical two-factor (species), two-level (inoculum density) design with center point. This hypothetical experimental design yields community compositions in which the growth responses of both species 1 and 2 are not affected by the design levels: Species 1 due to lower than measurable growth, species 2 due to saturating overgrowth. This is qualitatively representative of ER and BL in the DTL 1. Since the responses are not measurably correlated with the design variables, a predictive model cannot be determined. However, we reason that if a species overgrew, we should decrease the inoculum density, and if it undergrew, we should increase the inoculum density. **b** We update the levels by a half "frameshift" of the design levels (i.e. shifting the range of setpoints by half of its value in the desired direction). In other words, the new design uses the extremum of the old design as its new center point (low if overgrowth, high if undergrowth). Half of the new design thus overlaps with the old design, and the new "center point" level can be used as an input to any regression model, without forcing the model to extrapolate beyond training data, because it is a high or low level in the previous design. The hypothetical updated design results in the species growth responses being correlated with design conditions, which is qualitatively representative of the results of DTL 2. This approach flexibly integrates both experimental intuition and model-guided design. If modeling resources were not available, we expect this approach could be used to conduct a design-test-learn cycle without computational modeling, though we also expect it would be less efficient than model-guided design.

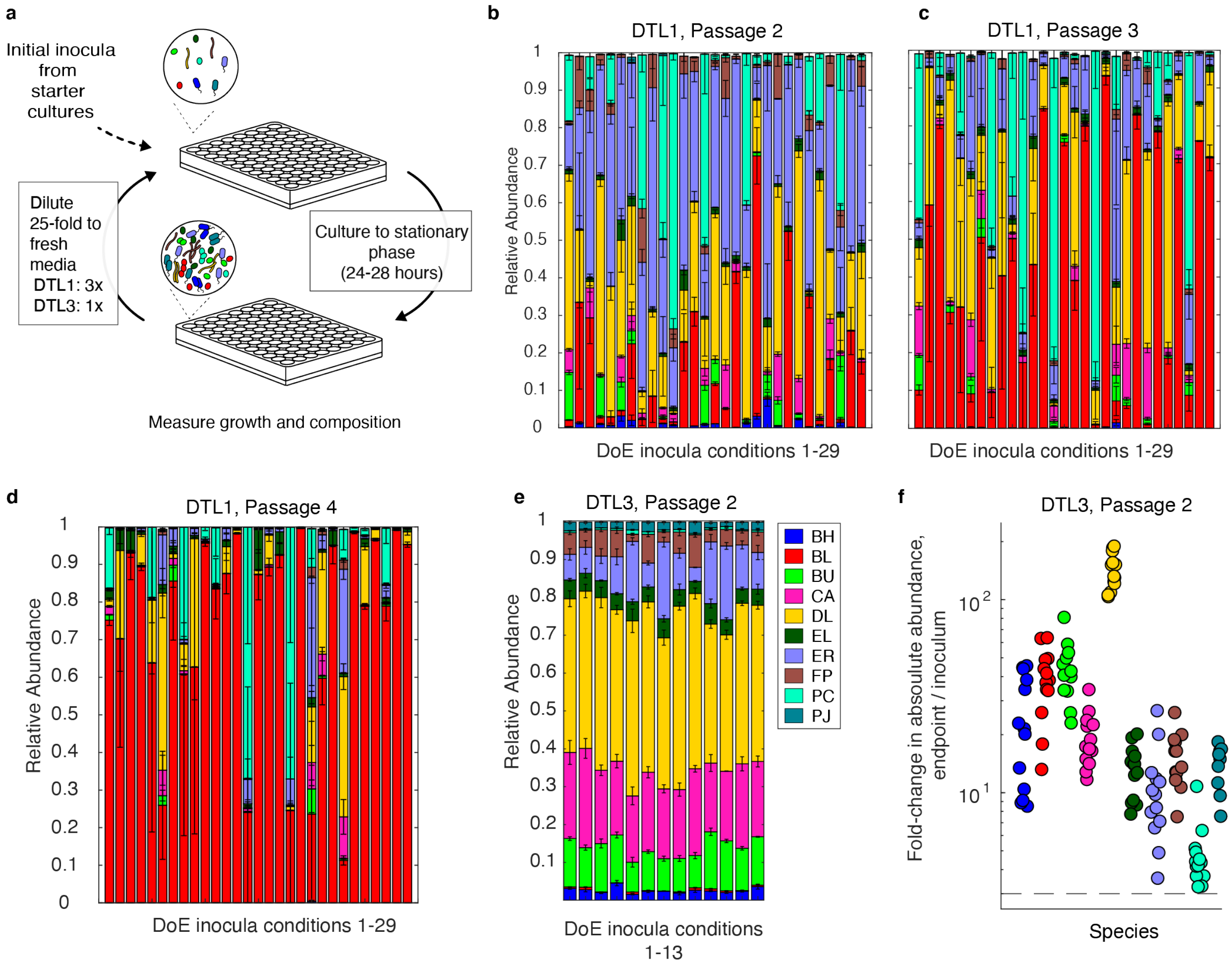


**Figure S9 Passaging training data for generalized Lotka-Volterra model training. a** Community passaging experiments are performed by diluting (25-fold volume/volume ratio) an aliquot of the stationary phase community cultures into fresh media, and then culturing these communities again until stationary phase. Measurements of community growth (OD600) and pellets for 16S rRNA gene next-generation sequencing are collected in the stationary phase of each passage. Stacked bar plots show the relative abundance (bar height denotes mean, error bars 1 s.d. from the mean of three biological replicates) for the three additional passages of DTL 1 (**b-d**) and the one additional passage of DTL 3 (**e**). **f** Scatter plot of the fold-change in the endpoint absolute abundance of each species divided by its corresponding inoculum absolute abundance for DTL 3, passage 2. The endpoint abundance of each species was greater than 3-fold (denoted by dashed line) compared to their initial abundance.

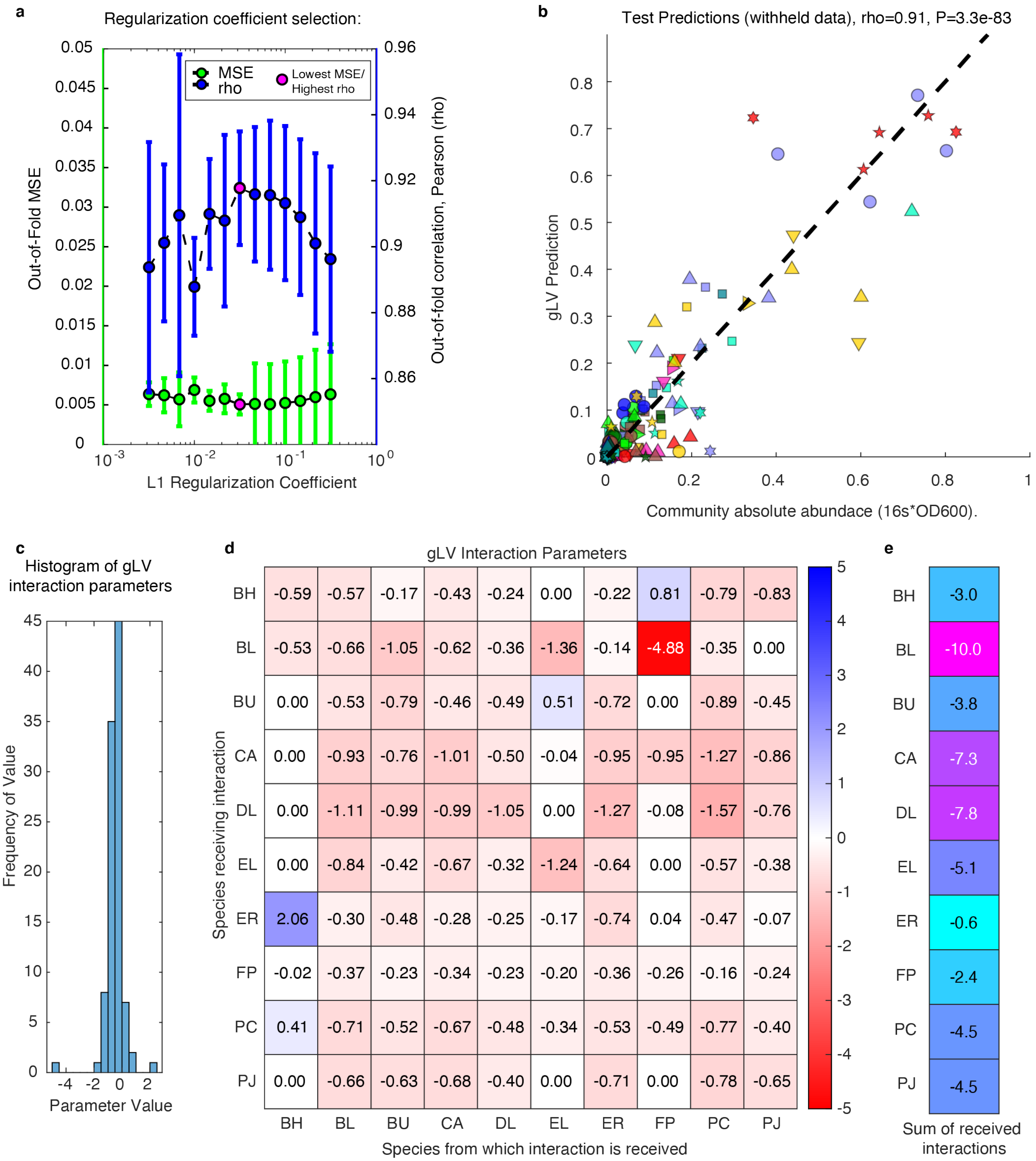


**Figure S10 Hyperparameter selection, predictions, and inferred parameters for the generalized Lotka-Volterra model. a** The L1 regularization coefficient is chosen as that which minimizes the average out-of-fold mean squared error (MSE) and maximizes the average out-of-fold Pearson correlation coefficient (rho) using 5-fold cross validation. Small regularization parameters lead to overfitting of training data and poor prediction of test data, while large regularization coefficients identify solutions that prioritize small parameter magnitudes over the quality of the model fit to the data. **b** Scatter plot of gLV model predictions on test data. Test data is randomly withheld from the entire fitting and hyperparameter selection process (using MATLAB's "randsample" function), such that the predictions on this dataset are an unbiased metric for model predictivity. Marker shape indicates training data experiment: circle – DTL 1, square – DTL 2, triangle - passage 2 of DTL

1, downward triangle - passage 3 of DTL 1, diamond – DTL 3, 5-sided star - passage 4 of DTL 1, 6-sided star - passage 5 of DTL 1, and right-pointing triangle – passage 2 of DTL 3. Pearson correlation (rho) and p-value (P). **c** Histogram of parameter values shown in heatmap in panel d. Many parameters are zero or near-zero because of the L1 regularization penalty. **d** Heatmap of interspecies interaction terms. The model's largest parameter explains ER's tendency to grow better in a community than in monoculture as a positive interspecies interaction with BH, while the smallest (largest negative) contributes to BL's tendency to undergrow in communities. **e** Sum of received interactions for each species (i.e. sum across rows of heatmap, $\sum_j a_{ij}$).

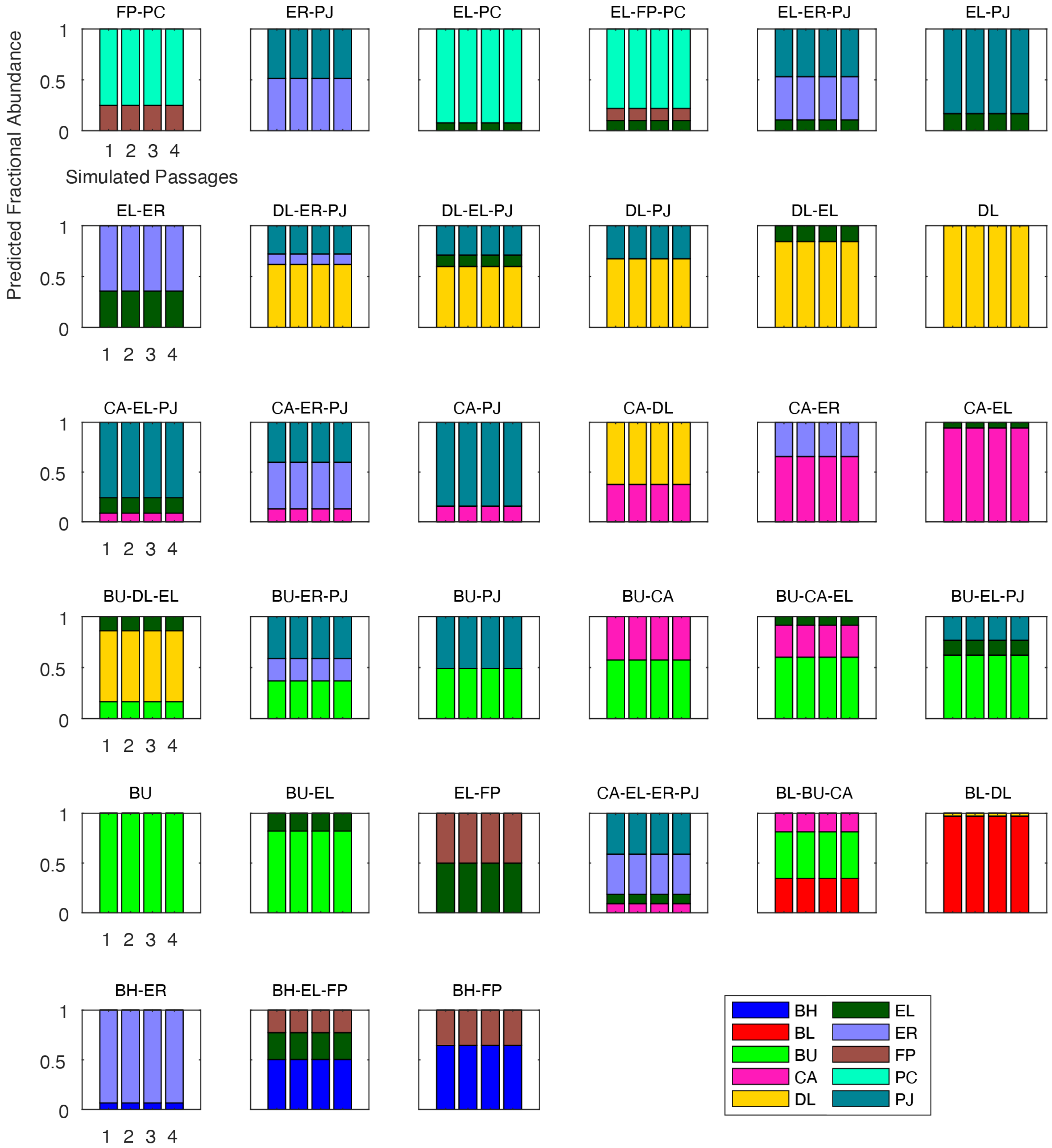


**Figure S11 The 33 unique solutions for low temporal variability of sub-communities based on optimization techniques.** Stacked bar plots indicate predicted species composition at stationary phase for four consecutive simulated passages. The initial conditions for each simulation are solutions to the low temporal variability objective function (equation 14). Of the 967 simulated subcommunities (10 choose k, k = 3 through 9", Methods), many converged to the same endpoint composition (within a numerical tolerance of 0.05 difference in absolute species abundance, as determined by MATLAB's "uniquetol" function, Methods).

**a**

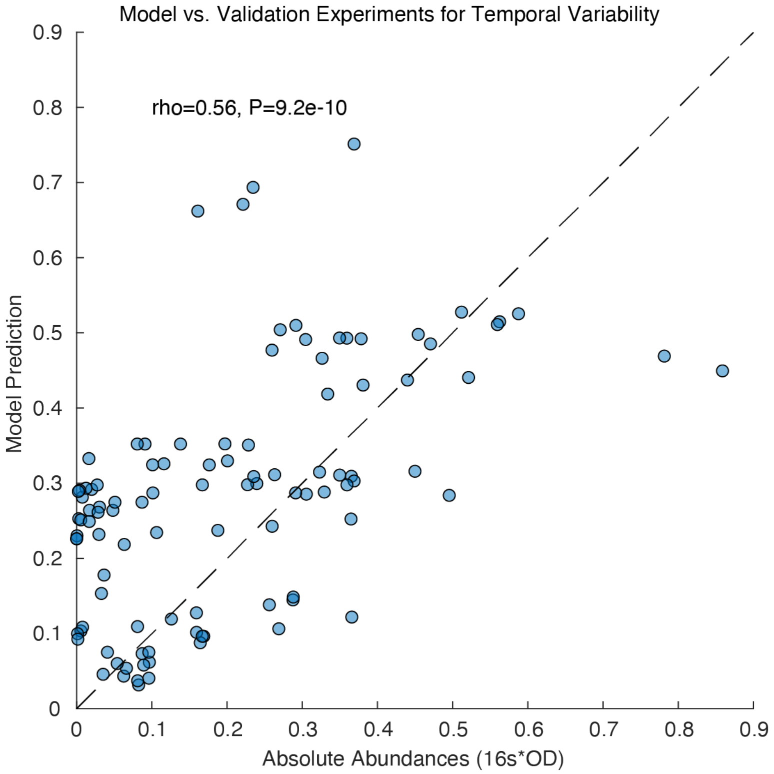


**Figure S12 Generalized Lotka-Volterra model predictive ability for low and high temporal variability designed sub-communities**. Scatter plot of species abundance predictions (low and high temporal variability community designs) vs. mean measured abundance in test communities (average of three biological replicates). The gLV model trained is trained on monoculture kinetic data, 10-member community initial conditions and passaging time points (3 passages from cultures in DTL 1 and 1 passage of the third). Validation measurements consist of a set of 2-4 member communities over four passages. Pearson correlation coefficient (rho) and p-value (P) are indicated.

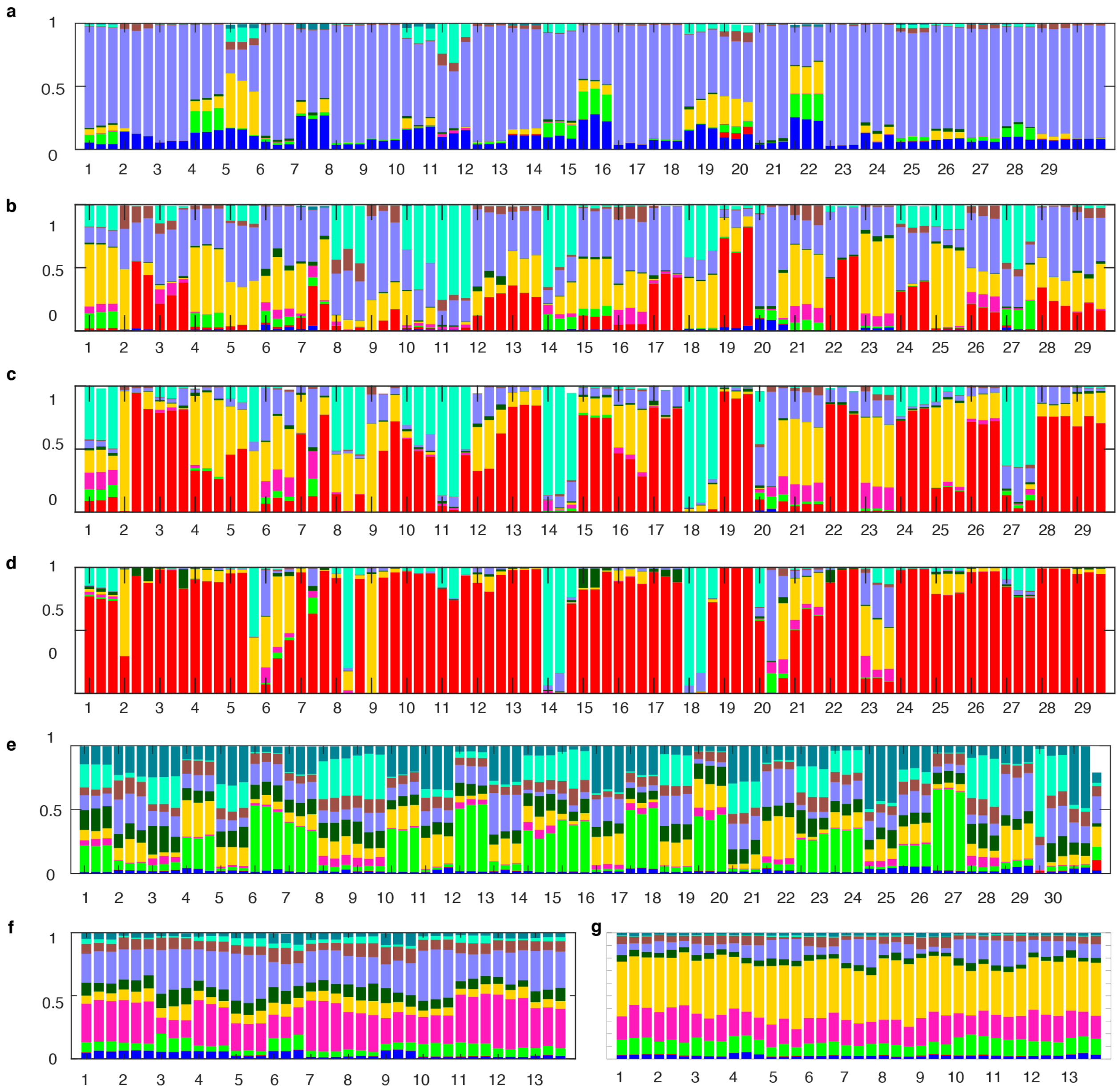

**Figure S13 Biological replicates for DTL cycle and passages.** Main text figures show mean and standard deviation of n=3 biological replicates to compactly visualize the many large datasets. Stacked bar plots show species compositions where colors indicate the relative abundance of each species based on 16S rRNA gene sequencing (y-axis) per corresponding main or supplementary figure, and x-axis has all replicates for each inoculation condition. The first replicate of each condition is labeled according to the DoE condition, second and third replicates are shown consecutively left to right. All biological replicates are shown as follows: **a** DTL cycles 1 (Fig. 3c DTL1), **b-d** DTL cycle 1 passages 2-4 (Fig. S9b-d), **e-f** DTL cycles 2-3 (Fig. 3c DTL2-3), **g** Fig. S9e.

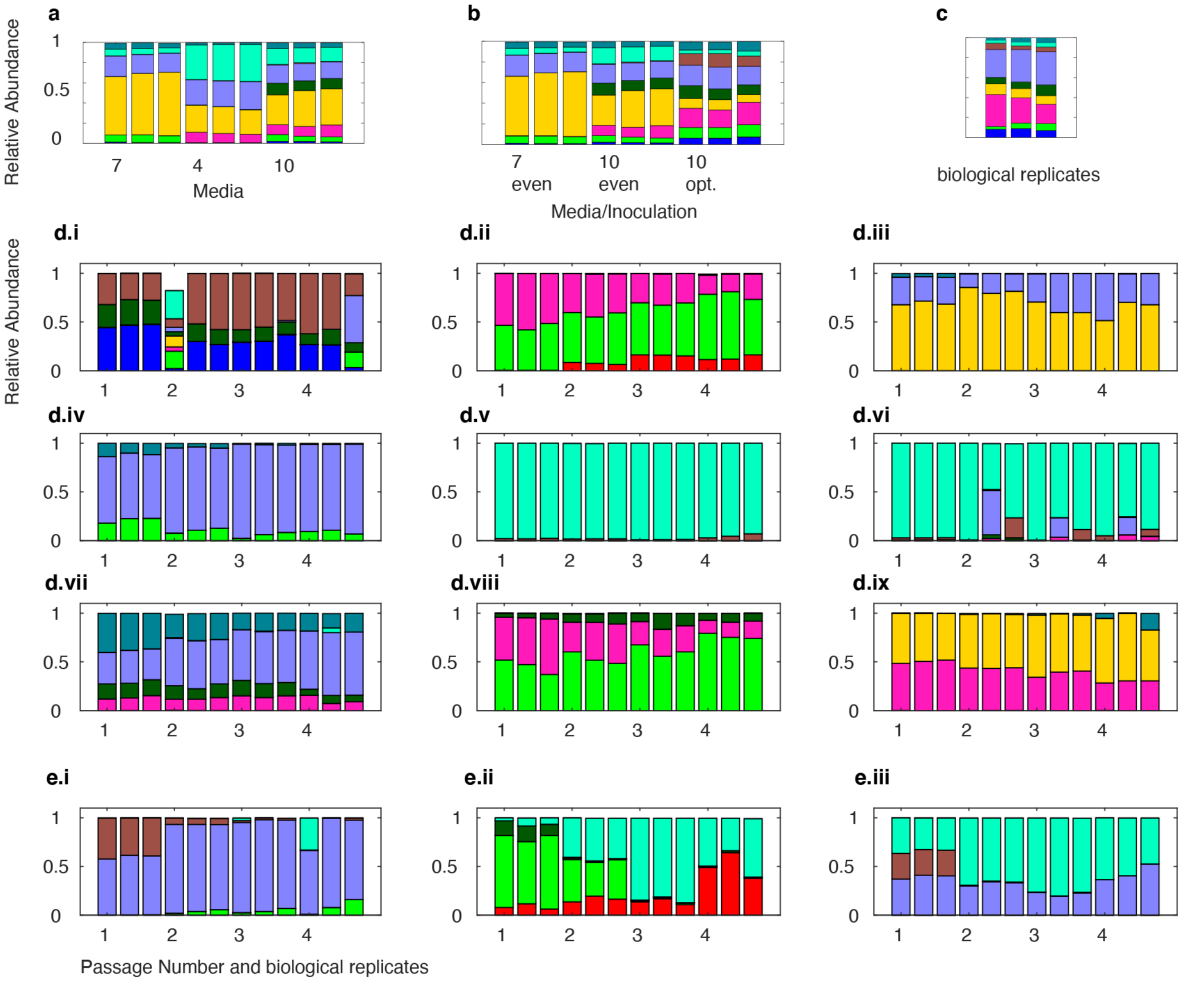


**Figure S14 Biological replicates for designed communities with high or low temporal variability.** Main text figures show mean and standard deviation of n=3 biological replicates to compactly visualize the many large datasets. Stacked bar plots show species compositions where colors species relative abundance based on 16S rRNA gene sequencing (y-axis) per corresponding main or supplementary figure, and x-axis has all replicates for each inoculation condition. All y-axes are scaled between 0 and 1. Biological replicates corresponding to **a** Fig. 1h where x-axis labels denote media type of first replicate, **b** Fig. 3i where x-axis labels denote media and inoculation conditions for first replicate, **c** Fig. 3h where x-axis shows biological replicates of 100mL scale up conditions (replicates of 200uL condition are shown in Fig. S13f condition 13), **d** Fig. 4d where x labels denote passage number of first replicate, and **e** Fig. 4c where x labels denote passage number of first replicate. Second and third replicates are shown sequentially left to right following first replicate. The following replicates were omitted from analysis (Methods) due to cross-contamination of >1% of total reads and/or low total sequencing reads <10% of average: **d.i** passage 2 replicate 1 and passage 4 replicate 3, **d.vi** replicate 2 passages 2-4.
